# DRUMS – A Flexible New Deep Learning Tool for Precise Cortical Surface Alignment across Individuals

**DOI:** 10.64898/2026.09.01.748559

**Authors:** Yubo Wang, Timothy S. Coalson, Chunhui Yang, David C. Van Essen, Matthew F. Glasser

**Affiliations:** Departments of Radiology, Washington University Medical School, St. Louis, MO, USA; Departments of Neuroscience, Washington University Medical School, St. Louis, MO, USA; Departments of Biomedical Engineering, Washington University Medical School, St. Louis, MO, USA

## Abstract

Accurate inter-individual alignment of human cerebral cortex is challenging because of the high variability of human cortical folding patterns and the regionally non-uniform and inconsistent spatial relationships across individuals of cortical folds versus the functional networks and cortical areas that we wish to study. To achieve precise alignment across individuals, an algorithm must use multi-modal neuroimaging features related to cortical areas and functional networks, ideally as inputs to surface-based registration, but at least during training if only folding patterns will be available during inference, so that it can learn where folding patterns are trustworthy and learn a spatially non-uniform regularization function that reflects the true gamut of human inter-individual variability in cortical organization. Additionally, an algorithm ideally will be capable of denoising its own input registration features to avoid overfitting to noise, enabling reproducible registration in test-retest data, and will not require precise hand tuning of input regularization parameters. To address these challenges, we developed Deep-learning Registration Using U-Net with Multimodal Supervision (DRUMS), a novel framework for multi-modal cortical surface registration. DRUMS applies its deep-learning approach to cortical surface registration using multi-resolution spheres, a well-validated approach used in other registration algorithms such as Multi-modal Surface Matching (MSM) (Robinson, et al., 2014; Robinson, et al., 2018). It also includes the same physically inspired strain energy regularization that we pioneered for MSM and the same precise barycentric interpolation on spherical surface meshes. DRUMS has a three-stage architecture: (1) multiscale feature extraction, (2) multiscale feature integration, and (3) deformation field generation. The framework’s multimodal design supports flexible integration of diverse imaging modalities, enabling registration using either folding features or multi-modal features as inputs with independent control over supervision during training (e.g., training DRUMS with folding inputs and multi-modal supervision to learn which folds best correlate with multi-modal features). DRUMS outperforms Multi-modal Surface Matching (MSM), the current state-of-the-art method used in the Human Connectome Project (HCP) pipelines, over a wide range of input regularization settings in both registration accuracy and test-retest reproducibility of registration results for both supervised folding registrations and multi-modal registrations across all tested modalities, including held out modalities. DRUMS further learns a biologically plausible spatially non-uniform regularization function, with registration induced distortion correctly matched to known human inter-individual cortical variability. These results suggest that DRUMS should replace MSM in the HCP Pipelines and position DRUMS as a versatile and reliable tool for cortical surface registration in neuroimaging research.

## 1 Introduction

In MRI-based neuroimaging, registration of individual brains to an atlas is a fundamental step in most preprocessing pipelines, enabling comparisons across individuals. Early efforts at MRI image registration used linear or nonlinear 3D volume-based methods (Ashburner & Friston, 2005; Jenkinson, et al., 2012). However, in humans such methods perform poorly in the highly convoluted cerebral cortex due to the extensive and regionally heterogenous individual variability of folding patterns and of cortical areal boundaries and functional regions relative to these folds (Glasser, et al., 2016; Coalson, et al., 2018). Surface registration (Fischl, et al., 1999; Van Essen, 2004) offers an attractive alternative approach to simplify the registration problem to 2D by ensuring that deformations respect the topology of the sheet-like cerebral cortex. However, surface registration works fundamentally differently than volume registration. Typically, the cortical surface is inflated to a sphere, and surface vertices are then deformed along the surface of the sphere so that their coordinates better match an atlas spherical mesh whose features occupy a specific geographic region on the sphere (e.g., the central sulcus spans a particular range of latitude and longitude values). The deformed individual surface and associated data can then be resampled to the standard mesh topology, thereby establishing vertex-wise correspondence between the individual and the atlas (or across individuals). Traditionally, folding patterns were used for surface registration (Fischl, et al., 1999; Van Essen, 2004); however, outside of early and primary cortices, many neocortical areas do not have consistent relationships with folds (Fischl, et al., 2008; Glasser, et al., 2016), and the folding patterns themselves often do not have a one-to-one relationship across individuals (Abdul-Kareem & TDCR, 2008; Leonard, et al., 1998).

These neuroanatomical realities have motivated the use of multi-modal surface registration to better align neocortical areas across individuals by using features that are more closely tied to cortical areas than cortical folds, such as myelination and resting state functional networks (Robinson, et al., 2014; Glasser, et al., 2016; Robinson, et al., 2018). Such approaches can markedly improve inter-individual alignment of human cortical areas (Coalson, et al., 2018); however, the method used for regularizing deformation is critical for optimally aligning functionally corresponding regions across individuals. Early methods of surface regularization in folding-based surface registration (Van Essen, 2004; Fischl, et al., 1999) tolerated extreme distortions of both surface area and surface shape and often did not optimize alignment of cortical areas, functional networks, or task activations, even when folding patterns became sharper in group averages, because their registrations overfit folding patterns relative to these cortical areal features of interest. Substantial improvements in folding-based alignment of cortical areas and brain function were achieved by highly regularizing the surface registration (Robinson, et al., 2014). This highly regularized folding-based surface registration approach, which was incorporated into the HCP Pipelines soon after their initial development, was termed Multi-modal Surface Matching (MSM) Sulcal registration (MSMSulc), and the regularization of the folding registration was carefully hand tuned to maximize multi-modal alignment, rather than folding alignment. However, if multi-modal features that are more closely tied to cortical areas like T1w/T2w myelin maps and resting state fMRI functional networks are directly available to a registration algorithm, less regularization is desirable to improve overall cortical alignment (termed MSMAll in the HCP Pipelines). Early MSM implementations (Robinson, et al., 2014; Glasser, et al., 2016) allowed the algorithm to “hide” distortion in certain unpenalized deformations. This limitation was addressed using a strain-based regularizer (Robinson, et al., 2018) that fully penalizes both areal distortion and shape distortion, enabling both more neuroanatomically plausible distortions and better interindividual alignment.

Although the strain-based MSMSulc and MSMAll alignments achieved state of the art performance for their time (Robinson, et al., 2018) and have been used by the HCP Pipelines and multiple HCP projects (Elam, et al., 2021) for many years, several limitations have become apparent. One important limitation is that each image registration is idiosyncratic, and the fine-scale results of the deformation are sensitive to noise (e.g., noise from the imaging process) in folding or multi-modal features, which means they are somewhat overfit to the data relative to hypothetical noise free data. This overfitting to noise in features reduces the test-retest reproducibility of the registration (e.g., evaluated using HCP participants who participated in two separate sets of scan sessions separated by weeks or months) and whose data is processed independently. Further, despite the improved behavior of the strain-based regularizer, the transition between reducing regularization enough to markedly improve alignment and reducing it so much that neurobiologically implausible extreme distortions occurred was relatively sharp and required careful hand tuning at the group level—and this threshold may not be the same for different individuals. In addition to differences across individuals, global regularization terms across the entire cortical surface may allow excessive below threshold deformations in many less variable regions while reducing the allowed deformations in regions that truly need the largest registration effects to fully optimize alignment. Additionally, some interindividual registration use cases (e.g., surfaces derived from post-mortem brain scans without any antemortem multi-modal MRI data) have only folding-based information, but the neocortical areal alignment performance of MSMSulc is less accurate than MSMAll (Coalson, et al., 2018). Because MSM is a classical optimization algorithm, it simply tries to best locally match the features it is given while minimizing the global regularization penalty, and it lacks high-level semantic representation of which features are reliable in different brain regions, how different brain regions differ in their individual variability, and how to distinguish noise from meaningful signal in the feature maps. As a result, MSM (like all other classical nonlinear image registration algorithms) overfits every registration it performs, leading to the limitations described above. Although MSM in principle could incorporate feature-specific spatially non-uniform cost function weighting and could apply spatial smoothing to the feature maps (though ideally such smoothing would also be spatially non-uniform), optimizing such parameters non-uniformly across the brain and appropriately for each individual using trial and error classical methods is impractical.

Deep-learning approaches offer potential solutions to all of the aforementioned problems (Cheng, et al., 2020; Li, et al., 2024; Suliman, et al., 2026). By learning registrations across a large group of individuals, rather than one at a time, deep learning approaches are less sensitive to the registration overfitting problem—to succeed, they must learn high-level semantic concepts that maximize alignment across an entire group of individuals, rather than focusing on idiosyncrasies of individual datasets that may not generalize. Additionally, deep learning approaches are effective at learning spatially non-uniform feature-specific cost function weightings (i.e., trust a particular feature in one location, but not in another) and spatial denoising filters. Such spatially non-uniform learning can substantially mitigate the tradeoff between regularization strength and alignment quality (versus a classical algorithm in which the regularizer must limit bad behavior everywhere and is set based on the worst location, even if lower regularization would benefit alignment quality across most of the brain), thereby turning the problem largely into a training issue (does training remain stable at a given regularization strength?). The group-wise training itself offers a strong regularization towards the correct solution, as overfitting noise or incurring excessive distortion in one individual will typically not be helpful in aligning others. Finally, deep learning approaches offer the opportunity to supervise training using features that will not be available at inference time (Li, et al., 2024), thereby enabling an algorithm to learn which portions of some features are relevant to the alignment of others (such as which cortical folds are well correlated with brain function or neocortical areas).

Indeed, several prior deep learning registration frameworks show promise for cortical surface registration. For example, (Cheng, et al., 2020) showed improved performance of a folding-based deep learning algorithm relative to their baseline FreeSurfer algorithm. However, the lack of replication in this study of the benefits of MSMSulc over that same FreeSurfer algorithm reported in multiple prior studies (Robinson, et al., 2014; Robinson, et al., 2018; Coalson, et al., 2018) is puzzling and raises concerns about potential biases in the study’s evaluation metrics. Similarly, (Li, et al., 2024) demonstrated marked improvements in functional alignment with folding using the cross-feature supervision concept during training mentioned above. Importantly, neither of these methods uses the standard spherical approach typically used in cortical surface registration. Instead, they cut the sphere along a prime meridian and project it to a 2D “map”, enabling classical 2D deep learning algorithms to be applied. This imposes large distortions associated with the flattening process (overemphasizing the poles versus the equator) and requires compensation for the cut topology. These distortions also interact with the chosen interpolation method within the algorithm, substantially reducing spatial precision of features near the poles with the standard bilinear interpolation typically used in 2D Cartesian deep-learning methods. Finally, this algorithm (Li, et al., 2024) proposes separating functional and anatomical coordinates; however, such a choice would not be neurobiologically well motivated, as brain function and anatomy are inextricably linked in each individual (if a neurosurgeon resects the wrong gyrus, the patient may suffer a neurological deficit). Deep learning has also been applied to spherical surface registration (Suliman, et al., 2026), showing improvements over MSM in both folding-based and multi-modal registration; however, this algorithm does not use the strain regularizer developed for MSM after extensive trial and error to fully encode both areal and shape distortions, and it does not incorporate cross-feature supervision during training. Also, they apply bi-linear resampling during training which leads to reduced feature precision in the map as in the 2D Cartesian method above. Finally, the spherical deep learning framework they used treated edges on ico-spheres as directed edges, which introduced a limitation in that convolutions occur only in one direction, which makes it harder for the algorithm to learn the correct registration deformation.

Here we propose the Deep-learning Registration using U-Net with Multimodal Supervision (DRUMS) algorithm, a spherical deep learning registration method that uses a biomechanically inspired strain regularizer and precise barycentric interpolation. Like MSM, DRUMS uses multiple spherical mesh resolutions and achieves rotation-robustness using geometric convolutions, which in our implementation are bidirectional convolutions. We implemented several distinct registration approaches: DRUMS-Sulc, a folding only registration; DRUMS-SupSulc, a folding registration that uses multi-modal features such as T1w/T2w myelin maps, and resting state functional network maps to supervise training (Li, et al., 2024); and DRUMS-All, which uses multi-modal features (including folding - unlike MSMAll), during both training and inference. We evaluate these methods using alignment quality both in modalities used for training and a held-out modality (task-based fMRI activations in held out individuals) and compare them with three other approaches: FreeSurfer’s classic algorithm, hand-tuned strain-based MSMSulc, and hand-tuned strain-based MSMAll. We also evaluate their areal and shape distortion and their test-retest reproducibility (in held out individuals). We show marked improvements in the alignment quality of areal features relative to FreeSurfer, MSMSulc (particularly with the DRUMS-SupSulc approach), and of DRUMS-All versus MSMAll. We also show marked improvements in test-retest reproducibility of the registrations and that our algorithm has learned a neurobiologically reasonable pattern of areal and shape distortions. These benefits are evident over a wide range of distortion penalty strengths, showing that DRUMS is less sensitive to this parameter than MSM. These findings result in a recommendation to replace MSM with DRUMS for the HCP Pipelines going forward for both folding-based and multi-modal areal-feature-based surface registration.

## 2. Methods

In this section, we present the data and computational preliminaries and then describe the details of our model.

### 2.1. Theory

#### 2.1.1. Icosahedral spheres

In this study, all data points (features) were positioned on the vertices of icosahedral spheres (ico-spheres) (Fischl, et al., 1999). Traditionally, ico-spheres have been constructed recursively. In the typical recursive process, new vertices are iteratively introduced at the midpoints on the edges of the existing triangles. This subdivision splits each triangular face into four smaller triangles, and the new vertices are projected outward to a sphere preserving the initial radius.

However, recursive ico-spheres constructed in this way exhibit uneven vertex areas, where the vertex area is defined as one third of the sum of the areas of the triangles sharing a given vertex. To enable this issue to be addressed, we implemented “wb_command -surface-create-sphere” in Connectome Workbench to generate optimized non-recursive ico-spheres, whose vertex areas are more uniform. Rather than using a recursive strategy, we split each edge of an icosahedron into an integral number of segments and tile the interior of each icosahedron face. To further improve the consistency of vertex areas on the sphere, we first compute weights to linearly interpolate positions for the new vertices from the initial icosahedron vertices but then apply an empirically tuned nonlinear function to each weight independently, use the altered weights to construct the initial vertex coordinates, and then project them to the circumscribed sphere. This nonlinear function was optimized only once, and was used for all spheres, such that exact multiples of subdivisions share the coordinates of the lower resolution’s vertices (though the vertex numbering is not shared). For the ease of implementing convolution operators, we use a 40962-vertex icosphere created with this strategy (ico-6) as the starting sphere, which is the same number of vertices as a recursive level 6 icosphere.

#### 2.1.2. The Spherical Registration Problem

In general, spherical surface registration includes four steps (Figure 1): 1. Map the anatomical surface to a sphere. 2. Initialization by rigid registration. 3. Non-rigid registration. 4. Resample the anatomical surface and other data with the new registered sphere. Here, we focus on the non-rigid registration step. Let *S*_*feat*_ and *T*_*feat*_ denote feature maps defined on the subject and template sphere (described in section 2.2). Consider a nonlinear transformation between spheres, Φ_*θ*_: *S*^2^ → *S*^2^, that represents an alignment of the subject features and the corresponding template features. The transformation is evaluated at a set of sample points *C*_*samp*_ on the sphere and is optimized to minimize the cost function:

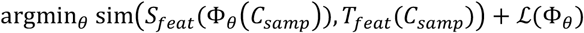

where sim(⋅,⋅) is a dissimilarity measure (described in section 2.5.1), and ℒ(Φ_*θ*_) is a regularization term (described in section 2.5.3) penalizing distortion. In practice, the sample points *C*_*samp*_ correspond to the vertices of the 40k-resolution icosahedral sphere described in Section 2.1.1. Both the subject and template spheres are represented using this common spherical mesh, providing a one-to-one correspondence between sample points and enabling direct comparison of feature values at matched spherical coordinates.

**Figure 1.**
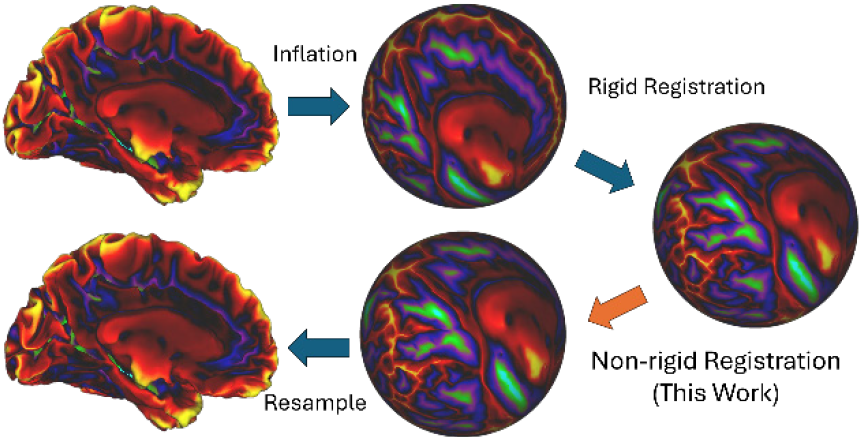
The process of surface registration.

#### 2.1.3. Resampling

Resampling is a critical step in spherical registration when applying a deformation field to align features between the two spheres. Traditional bilinear resampling, as employed by (Suliman, et al., 2026), projects spherical features onto a planar grid using longitudinal and latitudinal coordinates before performing interpolation. However, this projection introduces anisotropies in the precision of spatial features, especially near the poles, due to the distortion inherent in the coordinate transformation. To address this limitation, we adopted a more geometrically accurate barycentric resampling method, which operates directly on the spherical mesh, avoids such projection-induced artifacts, and better preserves mesh-local feature geometry across the sphere.

a. Barycentric resampling: Barycentric resampling entails two steps: vertex matching and resampling. Consider a vertex *t* on target mesh *T*. Vertex matching is performed by finding a set of vertices {*s*_1_, *s*_2_, *s*_3_} on the source mesh *S*, forming a triangular face which encloses *t*. Denoted the vector from vertex *v*_1_ to *v*_2_ as (*v*_1_ − *v*_2_). The sampling weight function 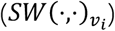 is defined as:

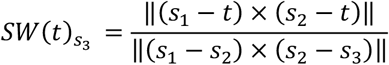

and similarly for the other two weights, where × is the vector cross product. The vertex *t* is projected onto the closest triangle in the source mesh, in which there exists a unique set of vertices {*s*_1_, *s*_2_, *s*_3_} that match with *t*. The resampling step can be considered as a weighted average using barycentric coordinates. Denoted *f*_*v*_ as the feature on vertex *v, f*_*t*_ is computed as a weighted sum:

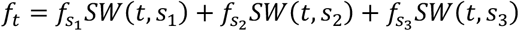
b. Adaptive barycentric area resampling (Glasser, et al., 2013) minimizes blurring during resampling while ensuring that all data from the source mesh contributes to the result on the target mesh. The method first computes both the barycentric weights (described above) of the target sphere vertices in the source sphere triangles (forward weights), and the barycentric weights of the source sphere vertices in the target sphere triangles (backward weights). For each vertex *t* on *T*, by definition, the forward barycentric mapping consists of the three vertices {*s*_1_, *s*_2_, *s*_3_} of a triangular face on *S* enclosing *t, F*(*t*) → {*s*_1_, *s*_2_, *s*_3_}. To establish a bi-directional matching, the roles of *S* and *T* are reversed. For each vertex *s* on *S*, the reverse barycentric mapping consists of the three vertices {*t*_1_, *t*_2_, *t*_3_} of a triangular face on *T* containing *s, B*(*s*) → {*t*_1_, *t*_2_, *t*_3_}. *B BF*(*t*) → {*s*_*i*_|*t* ∈ *B*(*s*_*i*_)}. This reverse barycentric mapping is rearranged to form a forward mapping, where each vertex *t* on *T* is associated with the set of source vertices {*s*_*t*_} that include *t* in their reverse barycentric mapping. These converted backward weights have an advantage when downsampling because there are as many weighting terms as for the equivalent upsampling, meaning that all vertices on the source sphere will contribute during downsampling, while simple barycentric downsampling can ignore source vertices that land in between vertices of the target sphere. For each vertex *t* on *T*, the forward mapping *F*(*t*) is used as the matching result if every element in *F*(*t*) is also present in *BF*(*t*). If this condition is not satisfied, the backward mapping *BF*(*t*) is used instead. We denote this new mapping as *M*(*t*). After generating these adaptive weights, the algorithm applies an area correction step using midthickness surfaces on the source and target meshes to derive the vertex areas. In this area correction step, the method multiplies the adaptive weights by the vertex area *VA*(*x*) of their target vertex *x*, normalizes them by the sum at their source vertices, then multiplies by the source vertex areas. For each vertex *t* on *T* and barycentric sampling weight *SW*(⋅,⋅), the corrected resampling weight is:

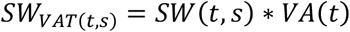

When an ROI map is provided, the source vertex is checked if it is inside the ROI. We then remove the vertex used in the resampling that was not in the ROI from the set *M*(*t*).

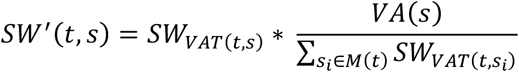

Finally, the feature *f*_*t*_ on vertex *t* is computed as a weighted sum:

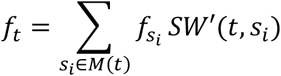

Although adaptive barycentric area resampling ensures the contribution of all input vertices, it introduces significant latency and increased optimization complexity during training due to its two barycentric computations and more complicated normalization. To mitigate these challenges, we instead adopted standard barycentric resampling during internal training steps, which does not have this guarantee but is computationally faster and made the network easier to optimize.

#### 2.1.4. Registration Drift and De-Drifting

Registration drift is defined as the group average registration effect, as the registration causes some areas to expand or shrink on average across the group. To evaluate the registrations fairly and to ensure that our registrations align to existing HCP MSM results, we introduced a de-drifting process that removed the drift between registrations. In this study, we de-drift all the registrations to MSMSulc, as MSMAll is also dedrifted to MSMSulc. We calculate the difference between the registered spheres from MSMSulc to the other registration, then we average such registration difference spheres across all the individuals. By applying the inverse of the average deformation to each individual’s registration using “wb_command -surface-sphere-project-unproject”, we eliminate the feature drift of registrations with respect to MSMSulc. All alignment and distortion metrics are computed after dedrifting. Notice that de-drifting is only performed once during the evaluation process and it will not be applied to new individuals.

### 2.2. Data

In this study, we utilized data from 1,071 subjects obtained from the WU-Minn Young Adult Human Connectome Project (HCP-YA) (Van Essen, et al., 2013). The HCP-YA collected and shared neuroimaging data from a large group of healthy young adults ages 22 – 35 (Elam, et al., 2021). HCP-YA datasets were collected from participants during a 2-day visit that included extensive behavioral testing plus four MRI imaging modalities (structural, resting-state fMRI, task-fMRI, and diffusion). MRI data were preprocessed using the HCP Pipelines (Glasser, et al., 2013) including structural preprocessing, functional preprocessing, and spatial-ICA FIX denoising and temporal ICA denoising (Salimi-Khorshidi, et al., 2014; Glasser, et al., 2018; Glasser, et al., 2019; Yang, et al., 2024).

For each individual, the dataset included a native cortical mesh for each hemisphere and the associated feature maps that were mapped to the individual’s native mesh. A 81-dimensional feature vector included one dimension for the FreeSurfer “sulc” map, one for the mean curvature map (Fischl, 2012), one for the folding-compensated cortical thickness map using nonlinear local multiple regression to reduce the effects of cortical folding on cortical thickness values (Demirci, et al., 2025), one for the T1w/T2w myelin map (Glasser & Van Essen, 2011; Glasser, et al., 2013), one dimension for an atlas level ROI (a mask for the non-cortical “medial wall”), and 76 dimensions for 76 components derived from resting state Probabilistic Functional Modes (PFMs) (Harrison, et al., 2015; Harrison, et al., 2020), which used hierarchical Bayesian models with priors for BOLD-like signal to identify resting-state networks (RSNs) without requiring either spatial or temporal orthogonality as in Independent Component Analysis (ICA) and performed joint group and individual inference. The details of how we organized data can be found in Section 8.1.

The 1071 individuals were split into three non-overlapping groups as described in (Yang, et al., 2025). One group of 475 individuals was used to train the network (training set), the second group of 475 individuals (with no family relationships to subjects in the first group) was used to evaluate the result (testing set), the third group of 50 individuals was used as a validation set for tuning the network parameters (validation set). 45 of the individuals in the testing set were scanned twice at an interval of several months to form a test-retest dataset, each having two versions of the surfaces and feature maps. These individuals were used to evaluate the reproducibility of the registration for different versions of the feature maps for the same individuals. Also, 40 of the 45 test-retest individuals were scanned with task-fMRI using an identical MRI protocol and preprocessing to the resting state fMRI while participants performed 7 tasks (working memory, gambling, motor, language, social cognition, relational processing and emotion processing), generating an additional 86 task feature maps (Barch, et al., 2013). These individuals were further used to evaluate the alignment of task features which were not used during training. The remaining 26 individuals were excluded to avoid family members being assigned across different groups to prevent genetic leakage (Yang, et al., 2025).

Training pre-processing includes three parts.

#### 2.2.1. Rigid registration and resampling

We first extracted the rigid rotation component from the standard FreeSurfer folding-based registration to fsaverage (Fischl, 2012) that was concatenated to the fsaverage to fs_LR registration (Van Essen, et al., 2011) by performing an affine regression operation on the native and fs_LR registered spheres. This process generated a rigid-registered native mesh sphere, which served as the initialization for the DRUMS registration (we have used a similar approach to initialize MSMSulc in the HCP Pipelines, where MSMSulc serves as the initialization to MSMAll). The registration features were then resampled from the rigidly rotated native mesh to the ico-6 sphere.

#### 2.2.2. Normalization and masking

The “medial wall” is a set of vertices needed to make the sphere a topologically closed mesh but that is not part of cerebral cortical grey matter. In the medial wall region, the “sulc” map and mean curvature map contain valid values, albeit not related to cortical grey matter features. The T1w/T2w myelin map values are not cortical, but they had a consistent pattern (primarily focused on the high T1w/T2w values of the white matter corpus callosum), and the cortical thickness values were constrained to be close to zero. Because of these consistencies, we elected to preserve the values in the medial wall region of these maps for supervision of alignment of the medial wall. For RSN values, we created a mask for “valid features” where the mask excludes RSN features within the medial wall while retaining all other features (as the fMRI features would be noise in this non-greymatter region). Using the features filtered by this mask, we performed normalization over the features (RSN features were normalized using the mean and standard deviation calculated from the valid region from all the 76 components). Finally, for the RSN channels, zeros were assigned to the medial wall regions. We found this approach produced more stable registrations around the medial wall than our prior MSMAll method that zeroed all features in the medial wall.

#### 2.2.3. Data augmentation

To improve the generalizability and robustness of the network, we introduced two types of augmentation during training.

a. Adding Gaussian Noise to Features: To enhance the network’s robustness to noise, Gaussian noise was added during training. Gaussian noise with *σ* was generated and added to the features. Subsequently, the mask described in the previous section was applied to reset the medial wall regions of the RSN channels to zero.
b. Adding Random Deformation to the Mesh: To improve the network’s robustness to individual variability among participants, random deformations were introduced to the mesh by adding Gaussian noise with *σ* to the vertex positions. To avoid highly focal distortions, the noise was generated at the ico-2, ico-3 and ico-4 levels, then up-sampled to the ico-6 level. Features were subsequently resampled based on the deformed mesh to the ico-6 sphere.

### 2.3. Template

The template used to train and evaluate the model was generated using 1071 subjects. It is necessary to start with an MSMAll aligned template because its features are substantially sharper than an MSMSulc template. Individual RSN maps on the 32k fs_LR mesh were projected back to the native mesh via the MSMAll registration. Then, all the features described in 2.2 were resampled to the ico-6 sphere using the MSMAll registered sphere and averaged to generate the template for registration.

### 2.4. Network

#### 2.4.1. Network Components

a. Gaussian mixture model convolutional (GMMConv) operator: Since the input features were positioned on a spherical surface, we applied the Gaussian mixture model convolution operator proposed in MoNet (Monti, et al., 2017). The key idea of GMMConv is to perform a weighted average of the features for a center vertex and all of its neighbors, then the resulting value will be assigned to the center vertex. There are two steps in this operator: 1. Transform the vertex coordinates from the Euclidean space to another space defined by pseudo-coordinates; 2. Perform patch-wise convolution.
  1. For a center vertex *a* of a neighborhood and each of its neighbors *b* on the sphere, we define the pseudo coordinate of the neighbor as:

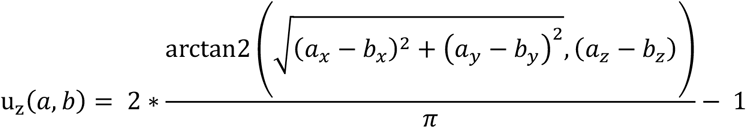

where *x, y, z* are the Cartesian coordinates and arctan2 is the inverse tangent. u_x_(*a, b*) and u_y_(*a, b*) are defined by analogy to form a 3-dimensional pseudo coordinate space.
  2. After transforming to the pseudo coordinate space, we then defined the convolution kernel using a Gaussian mixture model. Let *S*_*Θ*_(u) = (*w*_1_(u), …, *w*_*j*_ (u)) be the convolution kernel in which Θ was the set of parameters, u was the pseudo coordinate, *w*_1_, …, *w*_*j*_ were Gaussian kernels (before mixing). A single Gaussian kernel *w*_*j*_ was defined as:

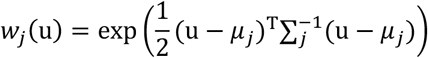

where *μ*_*j*_ was a mean vector and 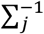 was a covariance matrix. For a 3d pseudo coordinate, *μ*_*j*_ had shape (3 × 1) and 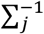 had shape (3 × 3). Next, we define the convolution operation. Let *f* be a function of feature maps where *f*(*x*) is the feature vector on vertex *x* and *g* be a Gaussian Mixing kernel,

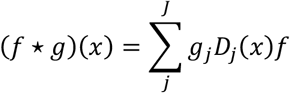

Further let N(*x*) be the neighbors of *x* (vertices with an edge directly connect to *x*), the patch operation was defined as

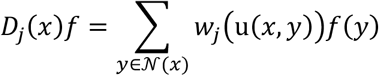

The detailed explanation of GMMConv can be found in Section 8.2.
b. Upsample operator: Based on the nature of ico-spheres from Connectome Workbench, we define upsample as the process of inserting new vertices into a lower-resolution ico-sphere (Figure 2). For each pair of adjacent vertices on the lower-resolution ico-sphere, a new vertex was introduced near their midpoint. Let *x* be the new vertex, *y*_1_ and *y*_2_ be the neighbors of *x* that are present in the lower-resolution sphere, the features at the new vertex were computed as the mean value of the features of the two adjacent vertices.

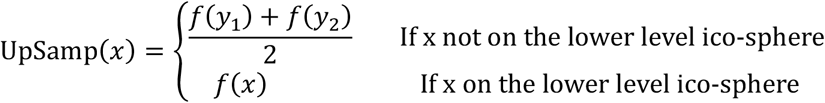
c. Downsample operator: We define downsample as the reverse of upsample (Figure 2). Given a higher-resolution source ico-sphere and a lower-resolution target ico-sphere, downsample involves reducing the vertex count by aggregating information. For each vertex on the target ico-sphere, there are six (or five, in specific cases) neighboring vertices on the source ico-sphere that are removed during downsample. The feature assigned to each vertex on the target ico-sphere is computed as the mean value of its own features and those of its neighboring vertices on the source ico-sphere.

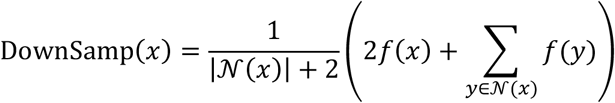

The reason for the 2*f*(*x*) term here is to compensate for the fact that the source vertices that exist on the target ico-sphere only contribute to one downsampled vertex, but the remaining source vertices contribute to two downsampled vertices.
d. Gaussian Error Linear Units (GeLU) activations: A variety of activation functions had been developed in recent years, but the Rectified Linear Unit (ReLU) style activation function remains widely used in convolutional networks due to its simplicity and efficiency (Liu, et al., 2022; Clevert, et al., 2016; Nair & Hinton, 2010). Here, we used GeLU activation function (Hendrycks & Gimpel, 2016), a smooth alternative to the tradtional ReLU activation function that uses Gaussian CDF-based smooth gating and has been used in BERT (Devlin, et al., 2019), GPT-2 (Radford, et al., 2019) and Vit (Dosovitskiy, et al., 2021). GeLU activation was defined as:

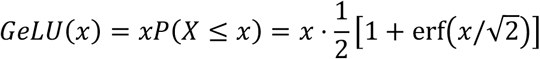

where erf is the error function. To improve the inference and backpropagation speed, this function was approximated as:

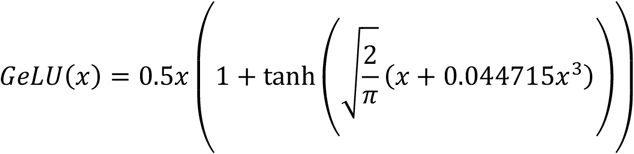
e. Batch Normalization (BN): Training Deep Neural Networks is complicated by the fact that the distribution of each layer’s inputs changes during training, as the parameters of the previous layers change. This slows down the training by requiring lower learning rates and careful parameter initialization and makes it notoriously hard to train models with saturating nonlinearities. To address this issue, we applied Batch Normalization (BN) (Ioffe & Szegedy, 2015) which performs the normalization and linear transformation for each mini-batch *x*:

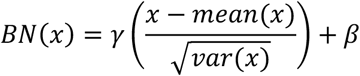

where *mean* and *var* are mean and variance functions, *γ* and *β* are learnable parameters for linear transformation.

**Figure 2.**
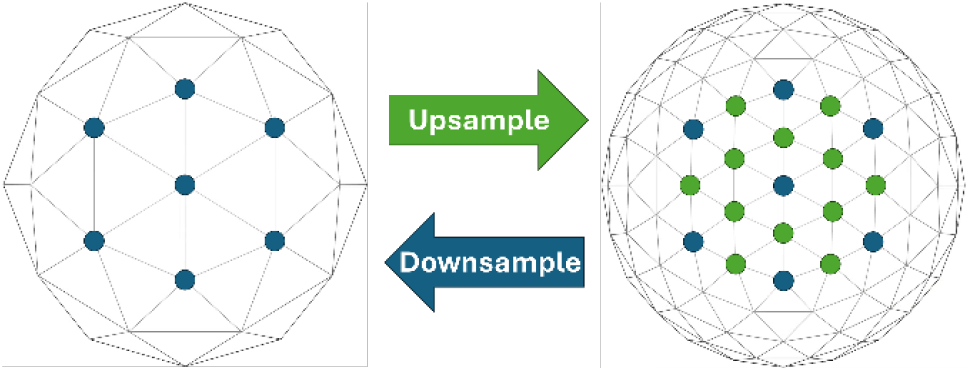
Demonstration of upsample operator and downsample operator.

#### 2.4.2. Network Architectures

a. Feature Transformation Block (FTB) (Figure 3.1): To efficiently mix the spatial-wise features and channel-wise features, we used a feature transformation block. Recent advances (Liu, et al., 2022) had demonstrated that the design principles of transformer-style networks can be effectively applied to convolutional networks to enhance performance. We incorporated several modifications to traditional convolutional network designs, including using fewer activation and normalization layers, GeLU activation. Based on the topology of the mesh, each convolution kernel operates on a vertex with its direct neighbors. To utilize the receptive field of such small convolution kernels, we stacked two convolution layers to achieve feature extraction for the first and second order neighbors.
b. Inverted Bottleneck (Figure 3.1.1): While convolution kernels are effective at extracting spatial features, the mixing of channel-wise information can be further improved. Inspired by recent architectures (Liu, et al., 2022; Sandler, et al., 2018), we incorporated an inverted bottleneck module that explicitly models interactions across feature channels. The module consists of two linear layers separated by a GeLU activation function. The first linear layer projects the features at each vertex into a higher-dimensional feature space, enabling the network to learn richer channel representations. GeLU activation introduces nonlinearity into the transformation, allowing more expressive feature interactions. The second linear layer then compresses the expanded features back to the original dimensionality, restoring the number of channels to match the module input. This expansion–nonlinearity–compression design enhances channel-wise feature learning while maintaining computational efficiency.
c. Attention Module (Figure 3.1.2): Recent research on attention mechanisms demonstrated that they can improve the representational power of a network (Woo, et al., 2018; Mnih, et al., 2014; Iandola, et al., 2016; Selvaraju, et al., 2017). An attention module enhances the networks’ ability to dynamically prioritize informative spatial regions or feature channels, thereby addressing a key limitation of traditional CNNs that treat all input regions uniformly. We incorporated an attention module inspired by the Convolutional Block Attention Module (Woo, et al., 2018) which assigns learnable weights to different parts of the input. There are two branches that process spatial- and channel-wise attention separately. In channel-wise attention, mean and max values across each channel are extracted and two linear transformations with learnable parameters and a non-linear GeLU function in between are performed separately to mean and max values, generating two channel vectors. A sigmoid function was then applied to the sum of the two channel vectors, and the resulted channel attention map was then multiplied by the original feature map. In spatial-wise attention, mean and max values across each vertex were extracted and convoluted using GMMConv, generating a spatial map with one channel. We then applied a Sigmoid function to the spatial map and multiplied it to the output feature map from channel-wise attention. By introducing such an attention module, the network handles input features with optimized spatial weights and further handles interactions between channels.
d. Encoder (Figure 4.1): Since the receptive field of the convolution kernel is fixed, we change the scale of the feature maps to improve the model’s ability to extract features at different scales (e.g., the convolution operation that covers the center vertex and its degree 1 neighbors on the 10k mesh covers a larger physical area than on the 40k mesh). We implemented a five-level U-Net encoder structure to extract and combine features across resolutions ranging from ico-6 (40,962 vertices) to ico-2 (162 vertices). At higher-resolution (lower) layers, convolutions captured local and detailed features, while at progressively lower-resolution layers, they extracted progressively higher-level global features. To ensure effective feature integration, skip connections (not shown) directly linked corresponding layers in the encoder and decoder, allowing the network to combine low-level detailed features with high-level semantic information (Zeiler & Fergus, 2013; Ronneberger, et al., 2015). The encoder has two input branches, one for a moving (deformed) image and one for a fixed (reference) image. The model will register the moving image to the fixed image. For individual to template registration, it registered an individual to the template, or for individual-to-individual registration, it registered individual1 to individual2.
e. Decoder (Figure 4.2): A single branch U-Net decoder was used to efficiently mix the features extracted from both moving and fixed images. The decoder first concatenated the output from the two branches in the encoder and further extracted/merged the feature maps from the ico-2 to the ico-6 scale. The result was then sent to a FTB and linear layer to generate the final deformation field.

**Figure 3.**
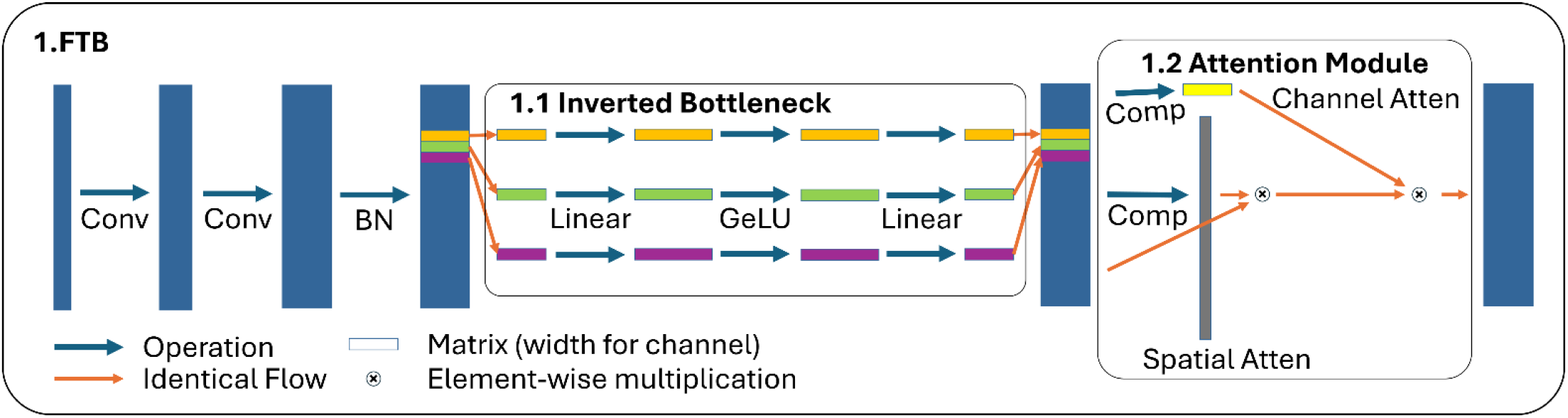
The Feature Transformation Block (FTB), Inverted Bottleneck module and Attention module. The blue and orange arrows represent operation and data flow respectively. The rectangles represent vertex-wise feature stored in matrix format. The height of the matrices represents the number of vertices, the width of them represents the number of channels. “Linear” operation means perform a linear transformation with learnable parameters across channels.

**Figure 4.**
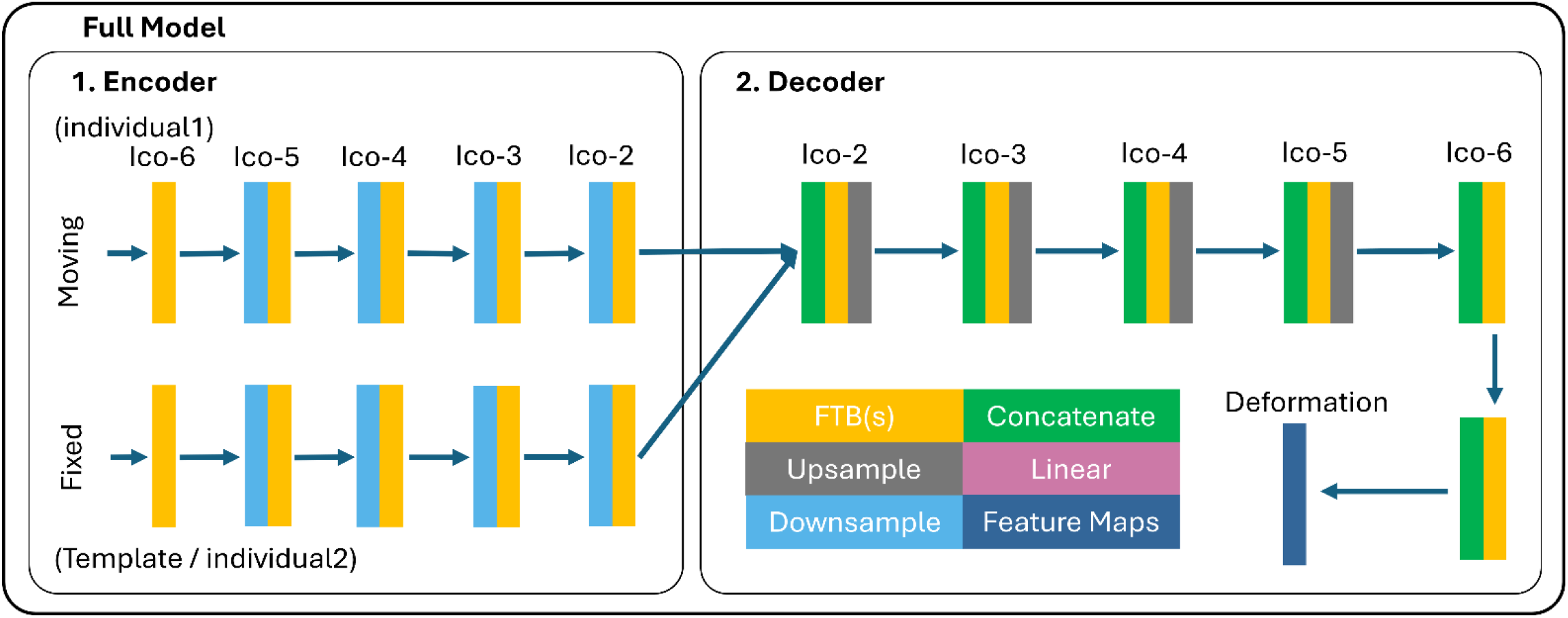
The DRUMS network. 1. Encoder: A U-Net style encoder with two branches that mixes the features for moving and fixed images with different scales. 2. Decoder: A U-Net style decoder that mixes the features from both moving and fixed images, generating the deformation field from those features. Each rectangle represents a specific operation (except the output deformation map), the arrows guide the direction of data flow.

### 2.5 Loss Functions and Evaluation Metrics

#### 2.5.1. Dissimilarity Loss

Features on the cortical surface typically exhibit significant differences between individual subjects and the template. We implemented cross-correlation (CC) as a similarity measure between the registered individual subjects and the reference. This measure accounts not only for pointwise feature differences but also for topological (spatial pattern) differences between the inputs after registration, enabling a more comprehensive comparison. The cross-correlation is defined as:

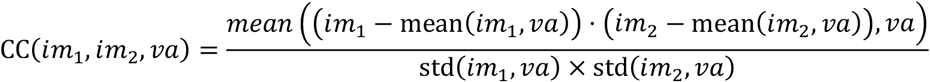

where *im*_1_ is the template, *im*_2_ is the individual subject being resampled using the registered mesh, *va* is the vertex area, ⋅ is the elementwise multiplication. We used weighted mean and standard deviation functions that used vertex area as a weighting factor to estimate cross-correlation on the anatomical surface. Since the gradient descent algorithms used to optimize the model minimize objective functions rather than maximize them, our model minimizes 1 − *CC* to improve similarity between subjects.

#### 2.5.2. Spherical Regularizer

We found that performing deformations in 3D Cartesian space (*x, y, z*) was more stable during training compared to 2D polar coordinates (*ϕ, θ*). To further enhance stability and ensure that the deformed mesh remains close to a spherical shape, we introduced a spherical regularizer that imposes additional constraints on the deformation. The spherical regularizer is defined as:

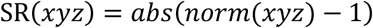

where *xyz* is a vector, norm is the L2-norm, and division is element-wise.

#### 2.5.3. Strain Regularizer (Figure 5)

We generalized the strain based regularizer in MSM and used it as a regularization during training. Consider for a triangular mesh face ℱ = {*p, q, v*} and the same face after deformation ℱ′ = {*p*′, *q*′, *v*′} where *p, q, v, p*′, *q*′, *v*′ are the positions of the vertex in a 3D space, there exists a local affine warp *F*_*pqr*_ that represents the 2D transformation matrix after projecting ℱ and ℱ′ onto the tangent plane. *F*_*pqr*_ fully describes the deformation of that triangle; its eigenvalues *λ*_1_, *λ*_2_ represent in-plane stretches. Then, the relative change in area can be described with *J* = *λ*_1_*λ*_2_ and the relative change in shape (aspect ratio) can be described with *R* = *λ*_1_/*λ*_2_. To penalize against both types of deformation, the strain energy density is defined as:

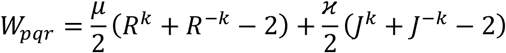

where *μ, ϰ* are hyperparameters to balance the terms. We set *k* = 2 to match the setting in MSM. To further constrain the pattern of distortion, we use the square of the sum of strain energy density across all triangular faces as our strain regularizer:

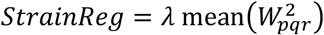

The hyperparameter *λ* balances the strength of regularization. A higher value of *λ* gives a stronger regularization to the model and reduces mesh distortions. We observed that DRUMS is more robust to the choice of *λ* than MSM, which allows for a wider range of distortions. The relation between amount of distortion and the alignment quality is discussed in 3.2.

**Figure 5.**
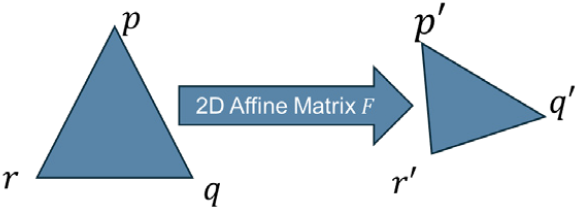
Graphical demonstration for strain regularization.

#### 2.5.4. Evaluation Metrics

Besides the similarity measure introduced in section 2.5.1, we also used cluster mass (CM) as introduced by (Robinson, et al., 2014; Robinson, et al., 2018) to evaluate the feature alignment. The cluster mass is defined as:

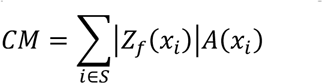

were *x*_*i*_ is a vertex coordinate, *Z*_*f*_(*x*_*i*_) is the z-transformed t-statistical value of the feature at this coordinate across registered feature maps of different subjects, *A*(*x*_*i*_) is the area associated with this vertex and *S* is a set of vertices where |*Z*_*f*_(*x*_*i*_)| > 5. In this case, A(x) is calculated from the midthickness surface registered with our model. Cluster mass is a more sensitive measure of changes in alignment quality than cross correlation, as it focuses on the alignment of peaks of activation in task fMRI GLM maps. Importantly, it is critical to resample both data and surfaces, and recompute group average vertex areas, before measuring cluster mass (or other evaluation metrics).

### 2.6. DRUMS Models

Previous studies showed that cortical folding patterns (sulc) are not well correlated with cortical areal features (e.g., myelin, RSNs, and task activation) across most of the human cerebral cortex (Glasser, et al., 2016; Coalson, et al., 2018). We implemented three versions of DRUMS to address different needs.

a. DRUMS-Sulc: DRUMS-Sulc registers the individual subjects to the template based only on the sulc and mean curvature features. It uses two input channels and generates a deformation field that maximizes alignment of the individual sulc and mean curvature map to the template. This method mainly provides a reference standard to compare the algorithm with other folding-based registration methods and to show the supervisory effect of DRUMS-SupSulc. It is not intended for general individual to atlas registration; however, we have found it useful for within-individual registrations of surface meshes generated from different datatypes (e.g., low-field MRI and tissue photo reconstruction), where folding patterns are expected to be corresponding, and it outperforms the highly regularized MSMSulc in aligning cortical folds in this setting when DRUMS-Sulc’s regularization is lowered.
b. DRUMS-SupSulc: DRUMS-SupSulc registers individual subjects to the template based on folding features. However, dissimilarity loss during training is supervised by areal features, enabling DRUMS-SupSulc to learn which folding features are reliable relative to areal features such as cortical myelin, thickness, and PFM-based fMRI measures. It uses sulc and mean curvature features as inputs but rewards or penalizes the algorithm based on whether the areal-feature-based correspondence improves or worsens. This method implicitly learns the coarse mapping from folding to areal features. We demonstrate below that it achieves higher alignment quality for functional features than DRUMS-Sulc or MSMSulc. Thus, it is our recommended initial folding-based alignment approach, replacing MSMSulc in the HCP Pipelines.
c. DRUMS-All: DRUMS-All registers the individual subjects to the template based on both folding and areal features. Unlike MSMAll, which is purely based on areal features, DRUMS-All uses sulc and mean curvature features as inputs to the network to stabilize the registration; however, their inclusion does not worsen the final functional alignment (and folding alignment is mildly improved relative to MSMAll). This method is our recommended multi-modal alignment-based approach, replacing MSMAll in the HCP Pipelines.

## 3. Results

We first present evaluation metrics and results for our model. We found that DRUMS-All and DRUMS-SupSulc tend to increase the relative size of regions having higher signal to noise ratio in their functional features and that this leads to registration drift, just as was previously shown for MSMAll (Glasser, et al., 2016). To compensate for this drift, all registration results were de-drifted to MSMSulc (as MSMAll was previously) to ensure a consistent alignment across methods. Attempts to incorporate anti-drifting penalties into the model did not improve alignment, though they increased model training time and complexity, so they were abandoned. We present two versions of DRUMS-All. The first uses *λ* = 0.03, resulting in a level of distortion comparable to that of MSMAll. The second uses *λ* = 0.01, allowing greater deformation to achieve superior alignment performance.

### 3.1. Alignment Quality and Amount of Distortion

In Figure 6 (left hemisphere) and Supplementary Figure 4 (right hemisphere), we present group average maps for each registration method. We arranged the methods into two groups based on their input features: folding-based methods (FreeSurfer, MSMSulc, DRUMS-Sulc, and DRUMS-SupSulc; columns 1 – 4 respectively) and those utilizing areal features (MSMAll and DRUMS-All; columns 5 - 7 respectively). Sharper group average patterns indicate more accurate alignment across subjects. To highlight prominent differences, we used arrows to identify regions with notable differences in local contrast between registration methods in the myelin (cyan and red arrows), RSN (cyan and green arrows), and task feature maps (yellow arrows). In these regions, DRUMS-SupSulc produced sharper and more spatially localized patterns than MSMSulc in the myelin, RSN, and task maps, indicating improved alignment of functionally relevant cortical areas. Similarly, both DRUMS-All settings generated sharper patterns than MSMAll across myelin, RSN, and task features, demonstrating superior alignment performance when multimodal features are used as inputs. To further illustrate the impact of registration at the individual level, we present RSN feature maps after registration in Supplementary Figure 2 and Supplementary Figure 3.

**Figure 6.**
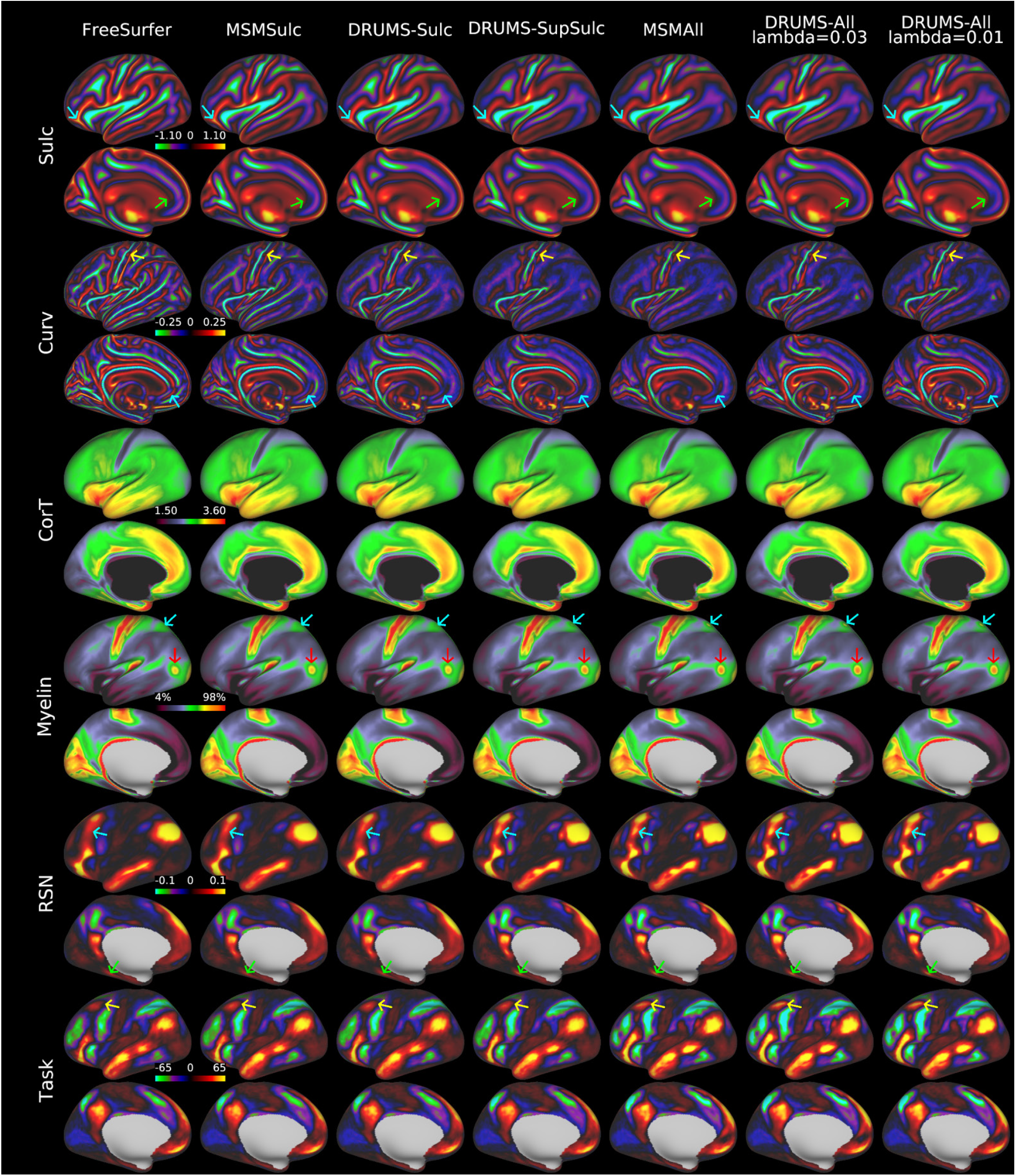
Group average across 475 subjects in the testing set. Each pair of rows show lateral and medial views of the left hemisphere. Each column shows a registration. There is no arrow for CorT row since the visual difference is not obvious. https://balsa.wustl.edu/VPDK7

Figure 7 (left hemisphere) and Supplementary Figure 5 (right hemisphere) show average areal distortion (abs Strain J), average shape distortion (Strain R) and group level feature drift from the template (Strain J), where some regions expanded (positive J) and some of them shrunk (negative J). The patterns of all maps were similar to MSMSulc because all registrations are dedrifted to it. The group average absolute areal distortion and average shape distortion showed where the models chose to make larger or smaller distortions when aligning the maps.

**Figure 7.**
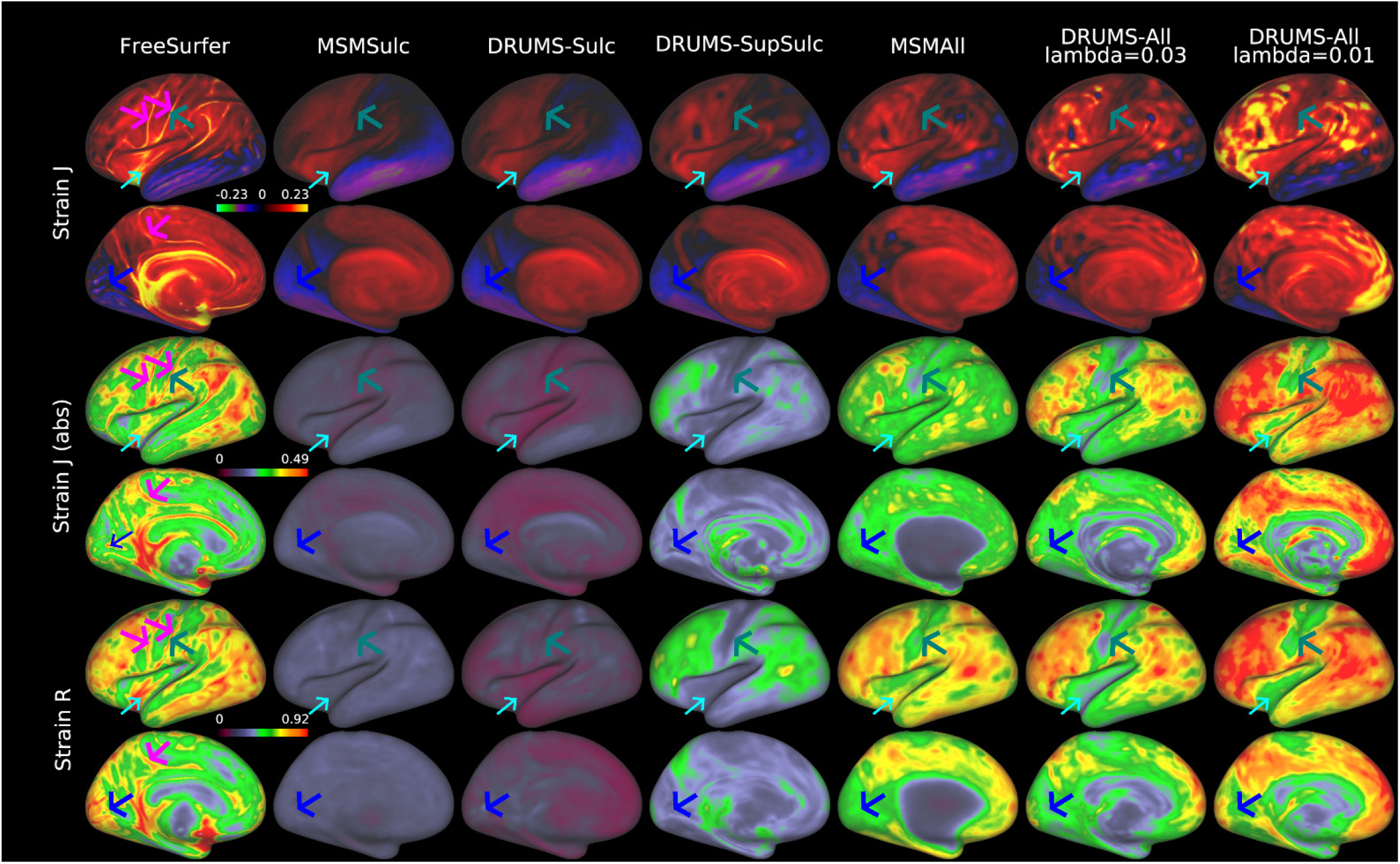
Group average distortion across 475 subjects in the testing set. Strain J and R values are log_2_-transformed. https://balsa.wustl.edu/80N2X

Importantly, the spatial pattern of distortions of DRUMS-All and DRUMS-SupSulc was neurobiologically plausible, as they match known patterns of individual variability in folding-areal feature agreement, with relatively less distortion around the central sulcus (teal arrows), the insula (cyan arrows), the calcarine sulcus (blue arrows), and along the medial wall (Glasser, et al., 2016; Coalson, et al., 2018) and relatively more distortion in other cortical regions known to have greater indivdual variability in folding-areal feature agreement (also seen in MSMAll to a somewhat lesser degree). These findings were particularly in contrast to the FreeSurfer alignment, which had increased distortion on gyral crowns (magenta arrows in the FreeSurfer column) relative to other locations and shows high distortion in regions of low individual variability. Overall, these results suggested that DRUMS successfully learned a spatially non-uniform biologically driven registration regularization function.

The qualitative observations were quantified in Table 1 (alignment quality) and Table 2 (amount of distortion). In Table 1, we used cross-correlation to measure the alignment of sulc, mean curvature (Curv), cortical thickness corrected for folding effects (CorT), myelin, RSN, and task features. Cross-correlation, defined identically to that used for similarity measurements during training (2.5.1), was measured on all the features used during training and in held out task-fMRI data. Alignment quality was also measured by the cluster mass of task-fMRI data, which was not included during training. We calculated the cluster mass of each channel of task-fMRI features, summed across channels, averaged across left and right hemispheres. Similar to MSM (Robinson, et al., 2014; Robinson, et al., 2018; Glasser, et al., 2016), we chose the model having the highest Task CM as the final model for each type of alignment.

**Table 1:**
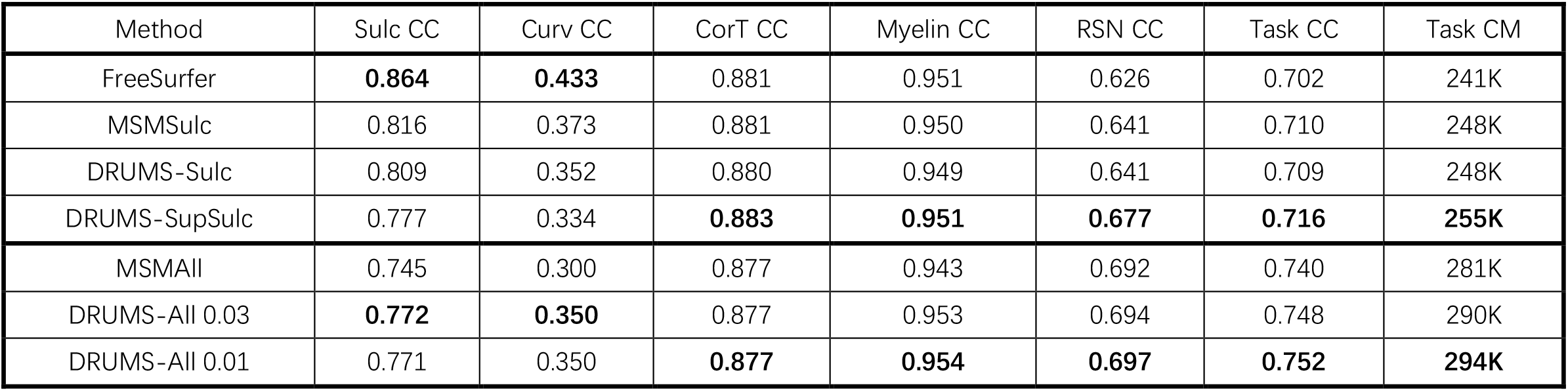
Comparison of alignment performance between FreeSurfer, MSM and DRUMS. Sulc CC, Curv CC, Myelin CC and RSN CC are calculated between individual and template used for training, Task CC is calculated between individual and group average. RSN CC is calculated using 76 PFM components (Harrison, et al., 2015; Harrison, et al., 2020) and taking the average. Task CC is calculated using 86 GLM maps (Barch, et al., 2013) and taking the average.

**Table 2:**
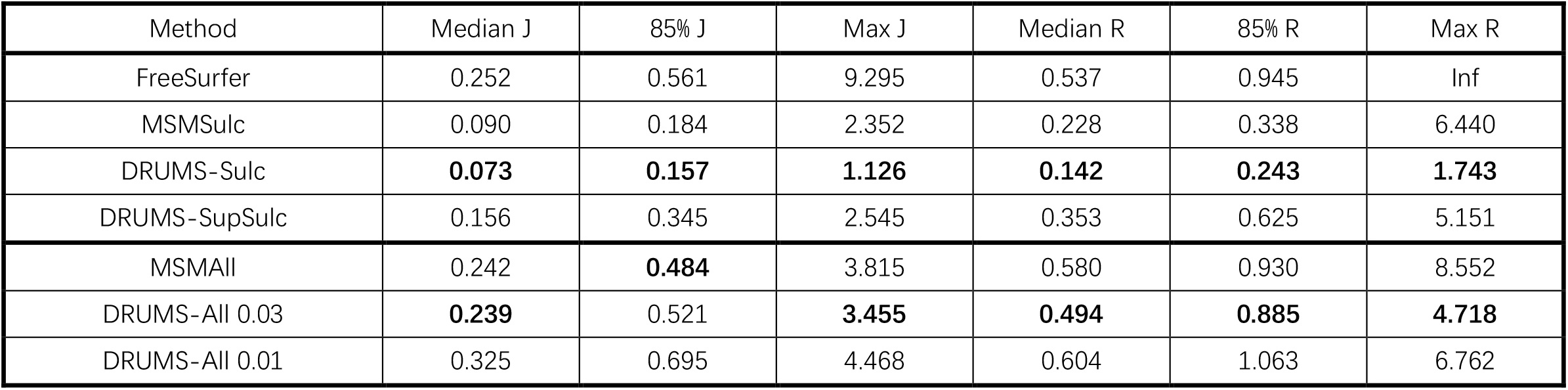
Comparison of distortion between FreeSurfer, MSM and DRUMS.

Table 2 reports the corresponding levels of distortion measured using strain J (areal distortion) and strain R (shape distortion). Log_2_-transformed values of J and R are reported to facilitate comparison across methods.

As shown in Table 1 and Table 2, FreeSurfer achieved the highest alignment scores for folding features. However, this improvement came at the cost of substantially poorer alignment of functional features, consistent with the blurred RSN and task patterns observed in Figure 6 and Supplementary Figure 4. Moreover, FreeSurfer introduced the highest levels of distortion among the evaluated folding-based registration methods.

Compared with MSMSulc, DRUMS-Sulc produced less distortion while achieving comparable alignment of functional features. Although its folding-feature alignment was slightly lower than that of MSMSulc at the regularization setting selected to maximize functional alignment, the two methods performed similarly on functional measures, indicating a more favorable balance between alignment quality and distortion.

DRUMS-SupSulc further improved functional alignment by incorporating multimodal supervision during training. It achieved the highest RSN alignment (0.677) and task cluster mass (255K) among registration methods that use only folding features as input. Although DRUMS-SupSulc introduced more distortion than DRUMS-Sulc, its overall distortion remained substantially lower than that of FreeSurfer while providing markedly improved alignment of myelin, RSN, and task features.

For areal-feature-based registration, DRUMS-All improved the evaluated alignment metrics relative to MSMAll. The DRUMS-All model with λ = 0.01 achieved the highest task cluster mass (294K) and substantially improved held-out task alignment at the cost of increased distortion. In contrast, the DRUMS-All model with λ = 0.03 maintained a level of distortion comparable to MSMAll while still providing superior alignment quality. These results demonstrate that DRUMS-All offers a more favorable alignment–distortion tradeoff than MSMAll across different regularization settings.

We further investigated the relationship between amount of areal distortion and task cluster mass caused by varying the registration lambda hyperparameter in DRUMS (explained in section 2.5.3). The values for MSMSulc and MSMAll were previously hand-tuned to be optimal; FreeSurfer’s is the default output of FreeSurfer’s recon-all.

In Figure 8, we present the alignment quality of task features under different amounts of distortion. The x-axis shows amount of distortion and y-axis shows task alignment quality, and demonstrates that DRUMS-All outperforms MSMAll and DRUMS-SupSulc outperforms MSMSulc over a wide range of lambda values. The results also highlight the substantial difference in performance between folding-based and areal-feature-based registration approaches. Although incorporating areal-feature supervision into the folding-based registration improved task feature alignment, a considerable performance gap remains between folding-based and areal-feature-based registration methods. According to the figure, task feature alignment improved for DRUMS-All as greater distortion was permitted under our experimental settings. However, we were unable to further increase the allowed distortion because training became unstable for *λ* < 0.009, and the minimal improvement on task alignment of *λ* = 0.009, compared to *λ* = 0.01 was not worth the markedly increased distortion suggesting that *λ* = 0.01 was near the end of the achievable alignment gains with the existing model design. Importantly, all of the tested DRUMS-All lambdas outperformed the hand-tuned MSMAll, illustrating a better alignment-distortion tradeoff of DRUMS-All relative to MSMAll with less than or a similar amount of allowable distortion; however, DRUMS-All was also able to further increase task cluster mass by reducing lambda and allowing more distortion than MSMAll before becoming unstable. DRUMS-SupSulc achieved maximum task alignment at *λ* = 0.05, with decreasing performance at lower lambdas. At this maximum level, functional alignment was clearly higher than MSMSulc or FreeSurfer. Like MSMSulc, DRUMS-Sulc performed best with very high regularization; decreasing regularization allowed overfitting of folding without improving functional alignment, analogous to FreeSurfer registration. Finally, it remained clear that even with DRUMS-SupSulc, where areal features are included in training to learn the reliable cortical folds, cortical folding remained unreliable across most of cortex, resulting in markedly worse performance of folding-based alignment relative to multi-modal areal-feature-based alignment (MSMAll or DRUMS-All).

**Figure 8.**
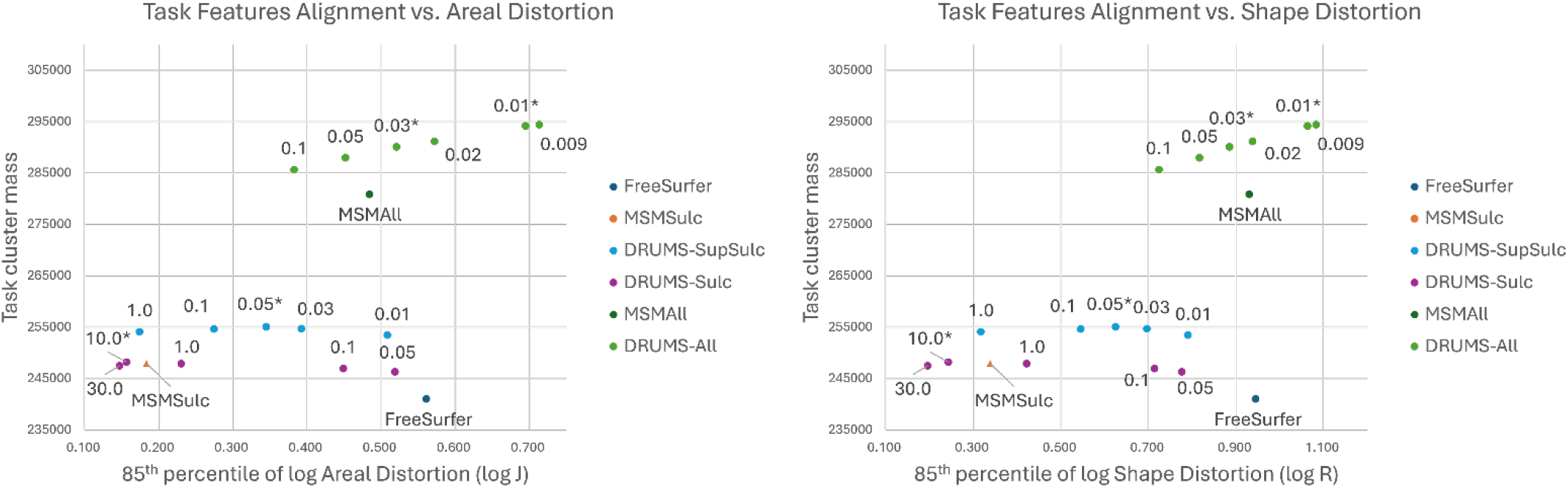
Relationship between task cluster mass and areal distortion. The x-axes show the 85^th^ percentile of areal distortion (left) and shape distortion (right) with log_2_ transformation and the y-axes show the task cluster mass. The triangles represent folding based registration; the points represent registration based on functional features. The lambda value used to regularize the models are labeled next to the data points (lower lambda allows more distortion). Notice that the lambda values are not comparable across different models due to their difference in loss functions. The results of models with lambda marked with * are reported in other tables and figures. We also present similar plots with median, 95^th^ percentile and maximum distortions in Supplementary Figure 7.

### 3.2 Test-Retest Reproducibility

Test-retest reproducibility is an important, but previously underrecognized, evaluation metric for registration algorithms. It provides insight into the sensitivity to local minima and idiosyncratic imaging noise (i.e., how much the resultant registrations are overfit to the provided data, rather than approaching the true optimal registration for the individual).

We evaluated the reproducibility of the midthickness surface to isolate the effects of the registration method while minimizing variability introduced by feature maps. For each subject, the midthickness surface was registered using different registration approaches for both test and retest datasets. An affine transformation from the test volume to the retest volume was estimated using boundary-based registration (Greve & Fischl, 2009) and subsequently applied to the test midthickness surface.

We then computed the Euclidean distances (*dist*3*d*) between corresponding coordinates on the midthickness surfaces for each test–retest pair. To further reduce surface misalignment arising from segmentation differences during cortical surface generation, we calculated the averaged signed distance (*distsigned*) from the test midthickness surface to the retest surface and vice versa, using the “wb_command -signed-distance-to-surface” function in Connectome Workbench. To approximate the tangential component of the distance between corresponding vertices, the final coordinate difference (*coord diff*) was computed as:

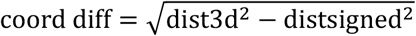

This formulation reduced test-retest variability introduced by inaccuracies in segmentation and surface placement, while preserving differences attributable to the registration model. Notably, it did not eliminate the influence of segmentation errors on the registration process itself (due to their effects on folding registration features), and thus on rigidly rotated registrations based on FreeSurfer.

In Figure 9 (left hemisphere) and Supplementary Figure 6 (right hemisphere), we observed a consistent spatial pattern of irreproducible regions (red arrow) across all registration methods around the medial wall, medial temporal lobe, insula, and temporal pole. We hypothesize that inaccuracies in brain tissue segmentation and surface placement lead to discrepancies in the reconstructed cortical surfaces between test–retest pairs. These surface inconsistencies propagated to both folding-based and functional feature maps, resulting in reduced reproducibility in the affected regions. Such variability was intrinsic to the upstream segmentation process and is therefore not readily mitigated by improvements in registration methods alone. Otherwise, it is clear that DRUMS-Sulc and DRUMS-SupSulc have smaller coordinate differences than MSMSulc and DRUMS-All has smaller coordinate differences than MSMAll. These findings are seen quantitatively in Table 3. The similarity (cross-correlation) between features from test-retest subject pairs are also shown in Table 3.

**Table 3:** Comparison of test-retest reproducibility between FreeSurfer, MSM and DRUMS. Similarities between test-retest feature maps are calculated using cross-correlation. Coord diff is calculated using mean squared distance between coordinates from the midthickness surface. The averaged result across all test-retest subject pairs is presented.

| Method | Sulc CC | Curv CC | CorT CC | Myelin CC | RSN CC | Task CC | Coord diff |
| --- | --- | --- | --- | --- | --- | --- | --- |
| FreeSurfer | **0.983** | 0.834 | **0.911** | 0.967 | 0.677 | 0.729 | 1.116 |
| MSMSulc | 0.976 | 0.803 | 0.906 | 0.964 | 0.675 | 0.727 | 1.287 |
| DRUMS-Sulc | 0.982 | **0.884** | 0.910 | 0.964 | 0.680 | 0.732 | **1.062** |
| DRUMS-SupSulc | 0.983 | 0.844 | 0.910 | **0.970** | **0.681** | **0.733** | 1.084 |
| MSMAll | 0.884 | 0.553 | 0.869 | 0.936 | 0.645 | 0.714 | 2.230 |
| DRUMS-All 0.03 | **0.972** | **0.785** | **0.910** | **0.969** | 0.693 | **0.731** | **1.405** |
| DRUMS-All 0.01 | 0.966 | 0.748 | 0.909 | 0.969 | **0.696** | 0.729 | 1.583 |

**Figure 9.**
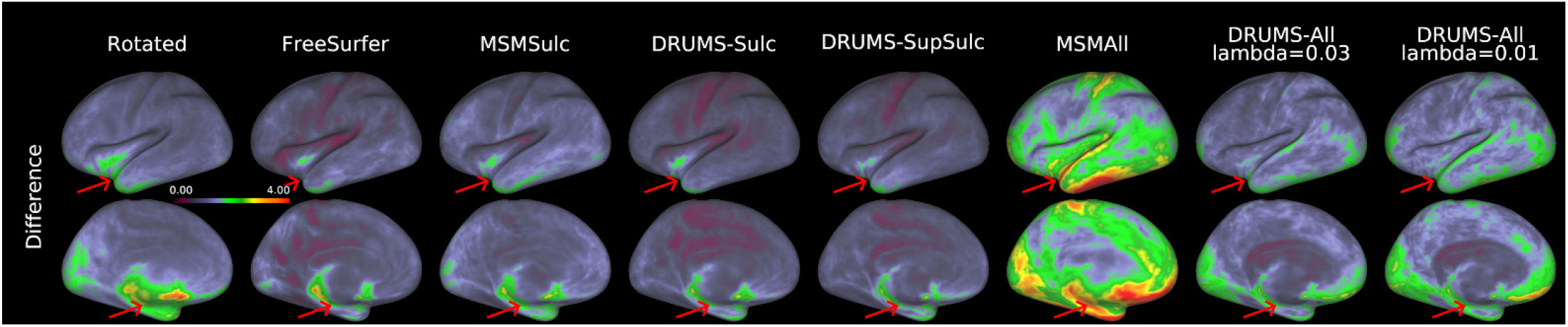
Differences in vertex coordinates between test–retest subject pairs for each registration method. Higher values indicate larger differences in vertex coordinates. https://balsa.wustl.edu/r6v77

As shown in Table 3, DRUMS-Sulc achieved test–retest reproducibility comparable to MSMSulc across all modalities. DRUMS-SupSulc further improved reproducibility for myelin, resting-state network (RSN), and task-based features. DRUMS-All achieved higher reproducibility than MSMAll, demonstrating that DRUMS produced reliable deformations and remained robust when used with noisier multimodal features. These findings match what we found in Figure 9 and Supplementary Figure 6. DRUMS-Sulc and DRUMS-SupSulc consistently outperformed both FreeSurfer and MSMSulc, while DRUMS-All consistently outperforms MSMAll under this evaluation metric.

MSMAll in particular had a weakness in test-retest reproducibility, as the relatively low regularization combined with a higher sensitivity of the classical registration algorithm to noise in the resting state network maps resulted in different test-retest registration solutions. Importantly, however, as noted above, DRUMS-All had much lower regularization and higher allowed distortion than all the classical registration methods, but maintained high test-retest performance. DRUMS-SupSulc showed a similar effect, but to a lesser extent. These results support our contention that deep learning registration can be relatively insensitive to noise, and as a result less likely to overfit in individuals, due to group-wise training, as discussed in the introduction.

### 3.3. Areal Classification

We further evaluated the performance of areal classifiers under different registration methods. The feature maps used by the areal classifier (Yang, et al., 2025) were resampled from their original MSMAll alignment to DRUMS-All alignment in one step. The areal classifier and subsequent post-processing pipeline were then rerun. Performance was evaluated using 45 test–retest subjects. For the 360 cortical areas (180 per hemisphere), we assessed classification consistency by comparing each parcellation result and summing the overlapping vertices. The mean consistency across all 45 subjects was then computed.

Alignment performance was evaluated using the following metrics:

a. Reproducibility between test–retest subjects, defined as the number of vertices assigned to the same cortical area in both test and retest sessions, which quantifies the stability of the parcellation.
b. Irreproducibility between test–retest subjects, defined as the number of vertices assigned to different cortical areas between test and retest sessions, which measures the number of vertices assigned to different areas across test–retest sessions.
c. Group overlap between test–retest subjects and the group-average parcellation, defined as the number of vertices for which both test–retest parcellations agree with the group-average result, which reflects agreement between individual parcellations and the group template.

As shown in Table 4, DRUMS-All achieves a higher group overlap compared to MSMAll, indicating a better alignment quality between individual subjects and the group average. It also achieves higher reproducibility and lower irreproducibility, indicating that DRUMS-All efficiently captures individual variability and is robust to the noise induced inconsistencies between test-retest subject pairs.

**Table 4:**
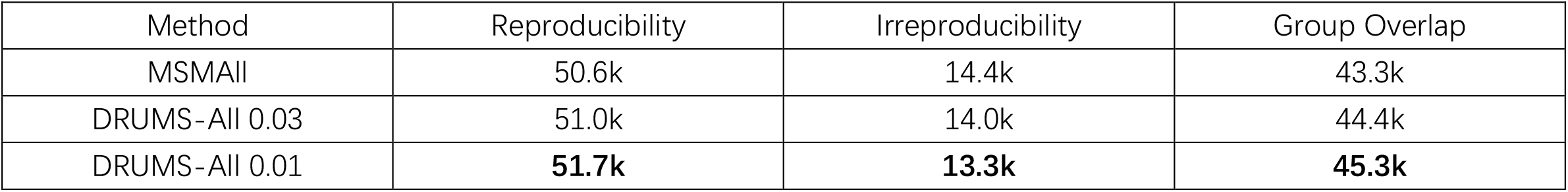
Evaluation metric for areal classification result. The results are the total vertex count that sum across 360 areas and both hemispheres then averaged across all the test-retest subjects.

### 3.4. “Medial Wall” Alignment

We also improved the alignment of cortex close to its boundary with the “medial wall” under functional supervision. In anterior cingulate cortex, there are many cases in which folding in the individual is poorly matched to the group average. In Figure 10, we presented an individual that has a double cingulate sulcus (red arrows) rather than the single cingulate sulcus in the group average (template column). Supervision by RSN and myelin (with medial wall regions masked off) introduces ambiguity in this region. We add supervision with cortical thickness and the full myelin map (along with adding the sulc and curvature channels), resulting in a reasonable deformation.

**Figure 10.**
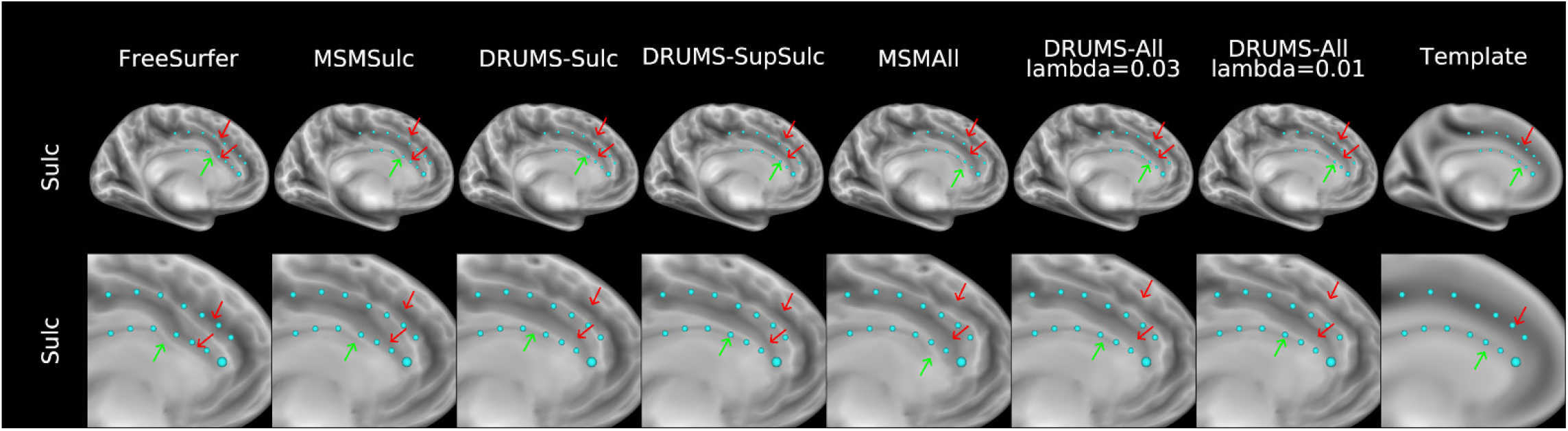
Demonstration of sulc alignment for a subject with a double cingulate sulcus. Each cyan ID node on the surface represents the same anatomical location. Ideally, the boundary of the medial wall should align with the inner set of the ID nodes. https://balsa.wustl.edu/nrnKk

In Figure 10, cyan ID nodes represent the same standard-mesh vertices across different registration methods, and green and red arrows show the key differences between them. We demonstrated DRUMS-SupSulc and DRUMS-All had the capability of correctly aligning folding features even with a different folding configuration. Compared to FreeSurfer and MSMAll, DRUMS models correctly aligned the boundary of the corpus callosum without being affected by folding variability of the cingulate sulcus.

## 4. Discussion

Here we present the DRUMS spherical deep-learning registration framework. It combines multiple key methodological advances including the use of deep learning on spherical surfaces, group-wise registration training, a strain-based regularizer shown to be useful in MSM because it fully accounts for registration induced distortion, improved interpolation methods during training and inference, and the use of areal feature supervision when training folding registrations. We show that DRUMS outperforms our prior standard methods of MSMSulc and MSMAll, which supports its replacement of these methods in the HCP Pipelines. The key advantages of DRUMS are the ability to learn feature-specific spatially non-uniform regularization penalties and the ability to ignore noise in the feature maps to both maximize alignment of areal features while also maximizing test-retest reproducibility. Indeed, DRUMS-SupSulc produces better alignment quality of areal features using folding features than MSMSulc (or FreeSurfer), indicating that it learns which folding features have reliable relationships with cortical areas across individuals and prioritizes aligning those, while otherwise minimizing distortion in other locations. DRUMS-All produces better alignment of areal features when areal features are available by enabling larger distortions in appropriate regions than MSMAll while at the same time preventing neurobiologically implausible extreme distortions or overfitting noisy feature maps that reduce test-retest reproducibility. Indeed, DRUMS-All learns a highly neurobiologically plausible pattern of distortions without user input, with regions known to be individually variable showing more distortions than regions known to be less variable, and it maintains high test-retest reproducibility even in the setting of higher distortions, indicating that it learns how to denoise feature maps as appropriate. Classical registration algorithms like MSM would only be capable of these properties with arduous hand tuning. MSMSulc and MSMAll were previously hand tuned with global regularization lambdas to maximize alignment of functional features while, in the case of MSMAll, maintaining a reasonable distribution of distortion, which already was an extensive neuroanatomist-driven process. Although MSM has the ability to use regionally specific regularization maps and could be implemented to have regional and feature-specific spatial smoothing, a deep-learning registration framework enables these properties to be learned automatically as a part of training while still having the advantage of group-wise training to prevent overfitting each registration to each individual.

DRUMS-SupSulc and DRUMS-All address different use cases, just as MSMSulc and MSMAll did. Folding-based alignment is important early in processing pipelines before multi-modal features are available to establish initial correspondence between an individual and a group average space. This correspondence enables generation of features like resting state network maps that rely on spatial embedding for their creation. (For example, a task GLM activation map can be computed in the absence of registration to a group and yet be matched across individuals through the temporal structure of the task, but resting state network maps cannot be generated in a corresponding way without a spatial mapping, as there is no temporal correspondence across individuals in the resting state.) Folding-based alignment is also important in situations where multi-modal features are simply not available. For example, the Human and Mammalian Brain Atlas (HMBA) project (The BRAIN Initiative Cell Atlas Network, 2026) is collecting deceased donor brains of individuals who died suddenly and whose families consented to donate their brains to enable a detailed understanding of the cell types and gene expression of human brain areas. These individuals are scanned using a very low field scanner producing a low resolution T2-weighted image that is used to create cortical surfaces after extensive post-processing (Gopinath, et al., 2026). DRUMS-SupSulc folding-based surface registration enables atlas human cortical areas from the HCP’s multi-modal parcellation to be projected probabilistically onto these brains as accurately as is currently feasible in the absence of antemortem multi-modal MRI. DRUMS-SupSulc enables both estimating the variability of each area after folding-based registration using this method (by generating a probabilistic map of each cortical area after DRUMS-SupSulc using the HCP-YA data where gold-standard individual parcellations are available) and then estimates the deformation to bring these probabilistic maps onto the individual’s cortical surface. Cell types and gene expression can then be measured in regions where there is reasonable confidence that a particular cortical area is located (Gao, et al., 2025).

DRUMS-All’s goal is to maximally align human brains where ante-mortem multi-modal features are available. At the same time, it is important to minimize overfitting to noisy feature maps such that the computed alignment is as close to the best-supported biologically plausible alignment. Such overfitting behavior will serve as nuisance variation when performing inter-individual comparisons for any group-wise analysis, a common goal of studies using the HCP Pipelines. Additionally, the individual variability of human cortical areas is of particular interest (Glasser, et al., 2016; Yang, et al., 2025); however, such analyses would be corrupted by registration-induced interindividual variability from overfitting to noise using MSMAll. Thus, more accurate estimates of the individual variability of human cortical areas will be produced by DRUMS-All than MSMAll.

Importantly, however, topology-preserving surface registration will always have a theoretical maximum performance. Although areal-feature-based topological variability appears considerably less than cortical folding variability, evidenced by the ability to produce much sharper group average maps of cortical myelin content, resting state networks, and task activation maps than folding maps (Glasser, et al., 2016), there is genuine topological variability in cortical areas (Glasser, et al., 2016; Yang, et al., 2025) and functional networks (Liu, et al., 2024). Fully characterizing such individual variability across the cortex remains an important topic for future work. Additionally, when group-wise analyses that take topological variability into account are desired, individual areal parcellation (Glasser, et al., 2016; Yang, et al., 2025), or functional network parcellation (Liu, et al., 2024) can be helpful. If a vertex-wise dense analysis is desired, methods such as non-topology preserving hyperalignment can be used instead (Haxby, et al., 2020). Importantly, all of these methods will be improved by starting from a highly accurate, topology-preserving alignment that compensates as effectively as possible for individual variability in the size, shape, and location of brain areas, networks, and functional regions.

In addition to accurately aligning features, DRUMS is also more convenient to use than MSM. DRUMS provides a flexible framework that allows users to easily modify input modalities and loss functions, enabling adaptation to different use cases such as non-uniform dissimilarity losses or customized distortion penalties. Computational efficiency is also substantially improved compared to MSM. For example, MSMSulc typically requires approximately one hour to register a single subject, and MSMAll may take several hours. In contrast, DRUMS requires only about 10 seconds per subject on a GPU and less than one minute on a CPU, significantly accelerating the processing pipeline. The training time is also within a practical range: DRUMS-All requires approximately two days of training on four NVIDIA A100 GPUs, while DRUMS-Sulc and DRUMS-SupSulc can be trained in less than 12 hours. These efficiency gains allow users to explore different configurations and tune the model within a relatively short period of time.

Another use-case for DRUMS in planned future work is inter-species registration (Hill, et al., 2010). The flexible DRUMS framework will enable multi-modal alignment of humans to macaques via Chimpanzees as an intermediate (Donahue, 2021). Although anatomical MSM (Robinson, et al., 2018) is potentially suitable for this problem, particularly if a spatially non-uniform differential penalization of areal and shape distortion has been implemented, so as to better account for different hypotheses of evolutionary changes in brain areas. However, using this tool would be an arduous semi-automated optimization process. A future, anatomically regularized DRUMS would be well positioned to automatically learn such mappings.

Although DRUMS presumably has limitations that will become apparent as it becomes more widely used, and it may require future refinement, it may be approaching the theoretical maximum performance of topology-preserving multi-modal image registration methods. In the history of cortical registration, moving from volumetric registration to surface registration was a major advance (Coalson, et al., 2018). Subsequently moving from folding-based registration to multi-modal registration was another major advance (Coalson, et al., 2018; Robinson, et al., 2014; Robinson, et al., 2018). Strain-based MSM over prior MSM methods was arguably a more modest advance, though virtuous in that alignment quality improved at the same time that distortion decreased (Robinson, et al., 2018). The gains from DRUMS-All over MSMAll are similarly of smaller magnitude than the folding vs multi-modal alignment gains, and this was the result of combining a large number of methodological advances. Although even smaller magnitude gains may be achievable, as topology preservation becomes less possible at progressively finer spatial scales of neuroanatomical organization, we anticipate that future advances will come primarily from non-topology preserving approaches and, potentially, from improvements in input modalities. Additionally, although there are efforts at integrating surface-based and volume-based registration (Postelnicu, et al., 2009), we suspect that such efforts will be challenged to overcome the fundamental limitation that folding patterns and where areas are relative to folds often simply do not correspond across individuals. As a result, such methods require extreme and neurobiologically implausible local distortions in the volume, such as turning a double Heschl’s gyrus into one. Indeed, rather than forcing the issue, we believe a better way forward is to use separate multi-modal alignment for surface-based and volume-based structures (Lange, et al., 2024).

## 5. Future work

Although additional small improvements may be possible by further parameter tuning, we instead propose two working directions that are related to computational geometry.

### 5.1. Creating uniform distributed pseudo coordinates

In this work, we use ico-spheres generated by Connectome Workbench for evenly distributed vertex surface areas on the sphere. However, *Figure 11* indicates that this distribution is not uniform in the pseudo coordinate space.

**Figure 11.**
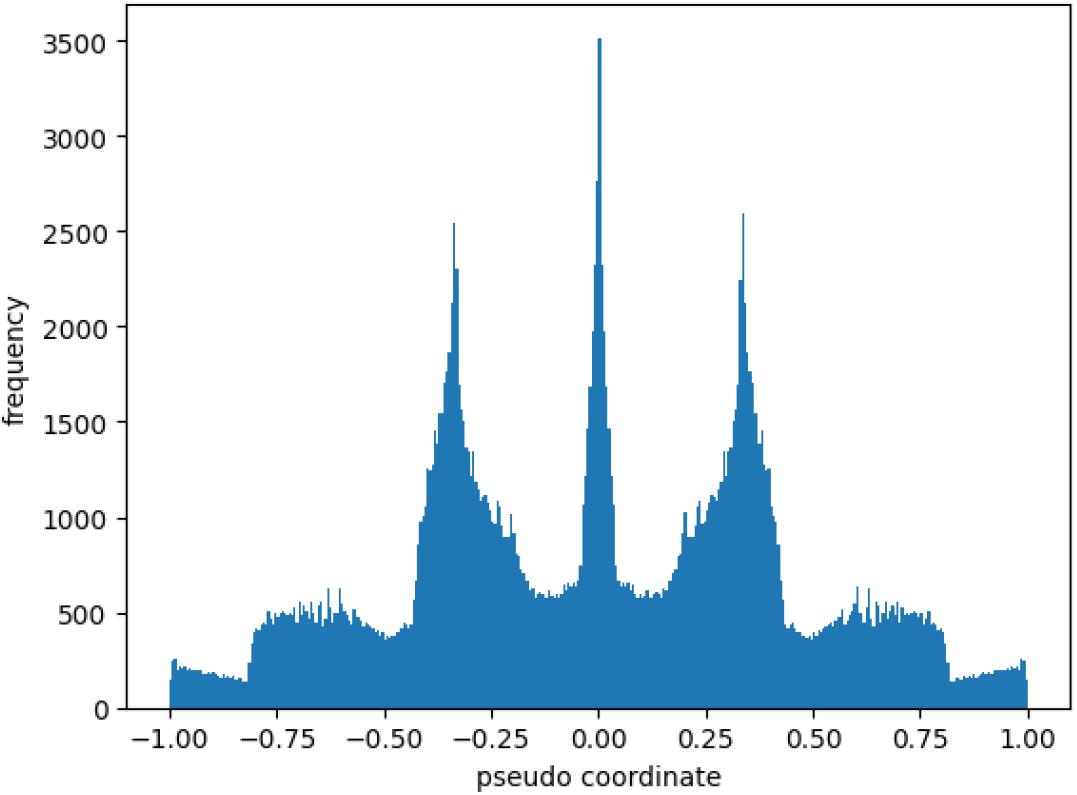
Distribution of pseudo coordinates representing angles between an edge and a Cartesian axis. The x-axis represents the pseudo coordinate that the edge is projected to, the y-axis represents the frequency of such pseudo coordinates.

The nonuniform distribution of pseudo-coordinates (section 2.4.1.a)) in the projected space may introduce bias in the learned convolution kernels. Constructing a new computational sphere or adopting an alternative projection method may help mitigate this bias.

### 5.2. Performing registration on the anatomical surface

In spherical registration, the anatomical surface is inflated to a sphere. However, this inflation introduces geometric distortions to the surface (Van Essen, 2004; Fischl, et al., 1999) that are not explicitly accounted for by the registration algorithm. The convolution operator used in the present study (Monti, et al., 2017) can in principle be applied to arbitrary surfaces, making it suitable for use on anatomical surfaces. To address this limitation, the spherical registration framework could be generalized to operate directly on the anatomical surface, with appropriate definitions for pseudo-coordinate projection and regularization.

### 5.3. Applying to more datasets

Applying DRUMS in other HCP-Style datasets is also desirable, including the HCP Lifespan and Connectomes Related to Human Disease datasets. However, DRUMS may not be suitable for non-HCP-Style datasets to the extent they do not meet important pre-requisites (e.g., fMRI denoising quality).

## 6. Conclusions

In this work, we introduce DRUMS, a novel deep learning framework for robust cortical surface registration. DRUMS captures individual variability and learns semantic feature maps, enabling accurate and reliable alignment. We generalize strain-based regularization into a flexible loss function, allowing for biologically plausible deformations. This formulation also enables users to control the regularization strength, offering a trade-off between alignment accuracy and deformation distortion.

DRUMS-SupSulc extends folding-based alignment by incorporating supervision from functional features, enabling cross-modality learning and multimodal supervision of a folding-input deformation model. Despite increased distortion, it substantially improves areal feature alignment compared to DRUMS-Sulc and MSMSulc.

With access to both areal and folding-based features, DRUMS-All achieves the best overall performance in aligning areal features, while maintaining a reasonable level of distortion. Additionally, it exhibits higher reproducibility, resulting in more reliable areal classification across test-retest sessions than MSMAll.

Overall, our results establish DRUMS as a powerful and flexible framework for cortical surface registration, setting a new performance baseline for HCP-style cortical surface registration under these metrics and motivating the switch of the HCP Pipelines from MSM to DRUMS.

## 8. Appendix

### 8.1. Data organization

In our framework, surface-related data were stored in GIFTI format, where feature maps and surface topology are represented separately for each hemisphere. Feature maps (e.g., sulc, myelin, RSN) are stored in GIFTI format (.shape.gii/.func.gii) as a matrix of size (*c, nv*) where *c* denotes the number of feature maps and *nv* denotes the number of vertices. Surface geometry is stored in a spherical surface file (.surf.gii) which contains two components: a vertex coordinate matrix of size (*nv*, 3) and a topology matrix of size (*nt*, 3) where *nt* is the number of triangular faces. The vertex coordinate matrix specifies the three-dimensional positions of the vertices on the sphere, while the topology matrix defines the mesh connectivity by storing, for each triangle, the indices of its three vertices. The ordering of vertices is consistent between the feature matrix and the vertex coordinate matrix, enabling feature values to be associated directly with their corresponding surface locations. The topology matrix then defines the neighborhood relationships among vertices used to construct the triangular mesh.

### 8.2. Detailed explanation of GMMConv

In this section, we provide a visual interpretation of the mathematical formulation of GMMConv. Supplementary Figure 1 illustrates the key steps of the convolution process. As shown in Supplementary Figure 1.1, features are represented as a matrix in which each row corresponds to a vertex and each column corresponds to a feature channel. We then apply the coordinate mapping described in Section 2.4.1.a.1, transforming the coordinates of neighboring vertices into pseudo-coordinates defined relative to the edge connecting each neighbor vertex to the center vertex. This transformation encodes the spatial relationship between the center vertex and its neighbors in a form suitable for convolution.

Supplementary Figure 1.2 provides an example of vertices after this coordinate transformation. The three axes represent the angles between the edge vector and the x-, y-, and z-axes in Euclidean space, respectively. As a result, each neighboring vertex is represented by a point in the pseudo-coordinate space. Gaussian mixture modeling is then performed in the pseudo-coordinate space, as illustrated in Supplementary Figure 1.3. Each Gaussian kernel assigns weights to neighboring vertices based on their pseudo-coordinates and produces a single output feature channel (shown in purple). Different kernels, initialized with different parameters, learn distinct weighting patterns and therefore extract different local features. Finally, the outputs from all kernels are concatenated to form the updated feature representation of the center vertex, as shown in Supplementary Figure 1.4. This process enables GMMConv to aggregate information from neighboring vertices while learning multiple complementary feature representations.

**Supplementary Figure 1:**
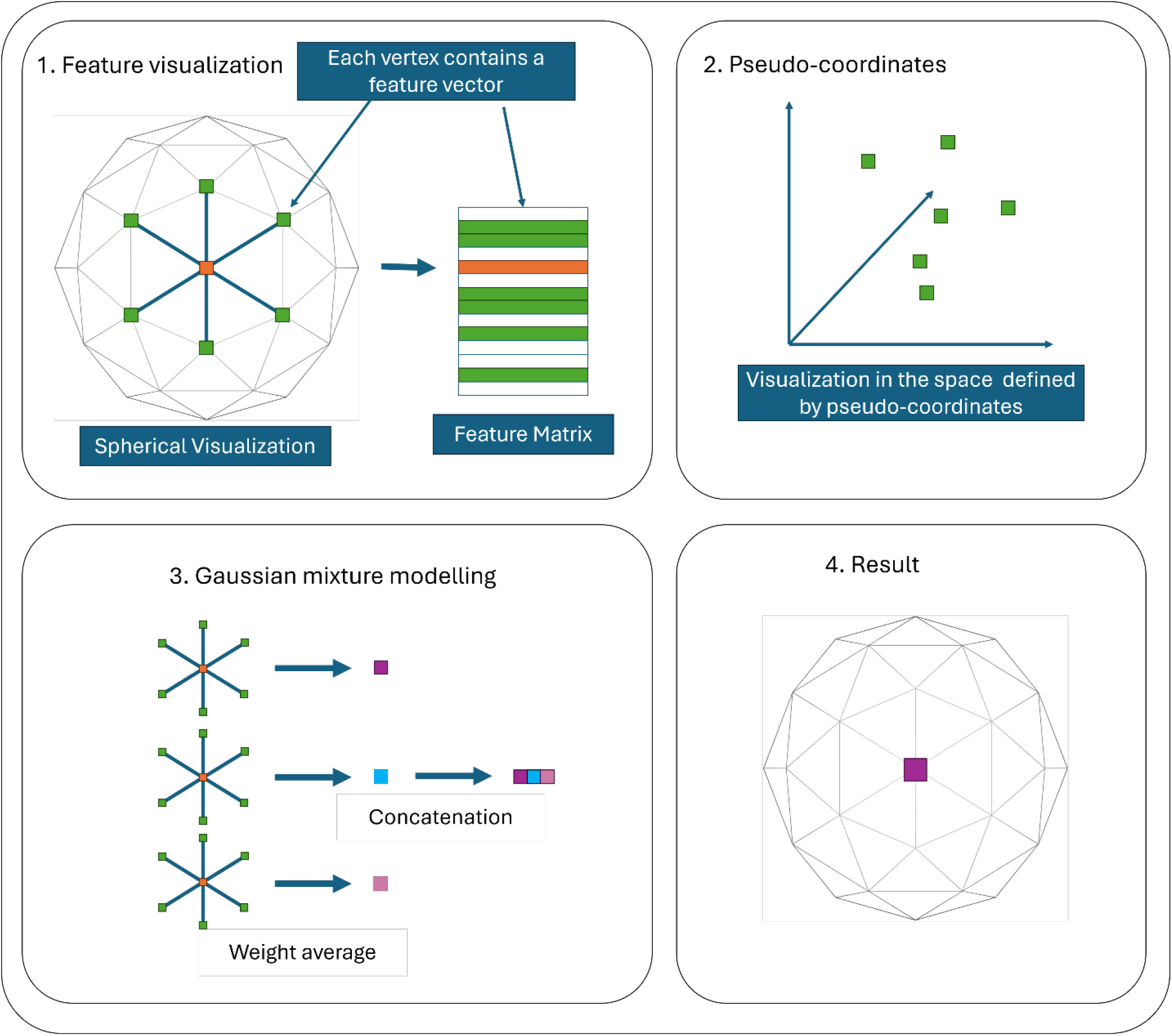
Visualization of GMMConv.

### 8.3. Training details

All models were trained under the ico-6 resolution. The U-net extracted features at scales from ico-6 to ico-2. AdamW (Loshchilov & Hutter, 2019) was used to optimize the model with 15 epochs of warmup. The exponential moving average (EMA) strategy was used to update the parameters, which was a variation of Polyak averaging (Polyak & Juditsky, 1992), because it stabilizes the training process and improves generalizability of the models.

#### 8.3.1. DRUMS-Sulc

The model took sulc and mean-curvature maps as inputs and was supervised by the cross-correlation of the sulc map. We added two extra feature channels that include sulc and mean-curvature around the medial wall for accurate alignment of that region. The weight was set to 10:5:1:1 for sulc, mean-curvature and the two extra channels respectively. We set the regularization weight to 10.0, learning rate to 1e-4 and channels for each resolution to (64, 96, 128, 192, 256). The model is trained for 500 epochs.

##### 8.3.2. DRUMS-SupSulc

The model took sulc and mean-curvature maps as inputs and was supervised by the cross-correlation of sulc, mean-curvature, cortical thickness, myelin and RSN features. We added two extra feature channels that include sulc and mean-curvature around the medial wall for accurate alignment of that region. The weight was set to 3:3:3:3:76:1:1 respectively. We set the regularization weight to 0.05, learning rate to 5e-4 and channels for each resolution to (64, 96, 128, 192, 256). The model is trained for 1500 epochs.

##### 8.3.3. DRUMS-All

The model took all the feature maps as inputs and was supervised by the cross-correlation of sulc, mean-curvature, cortical thickness, myelin and RSN features. We added two extra feature channels that include sulc and mean-curvature around the medial wall for accurate alignment in that region. The weight was set to 1:1:1:1:76:1:1 respectively. We set the regularization weight to 0.01, learning rate to 1e-4 and channels for each resolution to (64, 128, 192, 288, 432). The model is trained for 1500 epochs.

### 8.4. Example subjects for RSN alignment

For further understanding of how registration affects the alignment of individual subjects, we present the alignment of a PFM (Harrison, et al., 2015; Harrison, et al., 2020) component derived from resting state fMRI. We present the result from both multi-modal registration (MSMAll and DRUMS-All) and folding based registration (MSMSulc and DRUMS-SupSulc).

In Supplementary Figure 2, we present an example of PFM components derived from resting state fMRI for a single subject. We mark two high-intensity regions in the template RSN component. In this example, MSMAll misaligns both regions, with only approximately 30% of the regions overlapping with the template. In contrast, DRUMS-All achieves better alignment, with most of each region correctly aligned to the template, resulting in improved overall alignment quality.

**Supplementary Figure 2:**
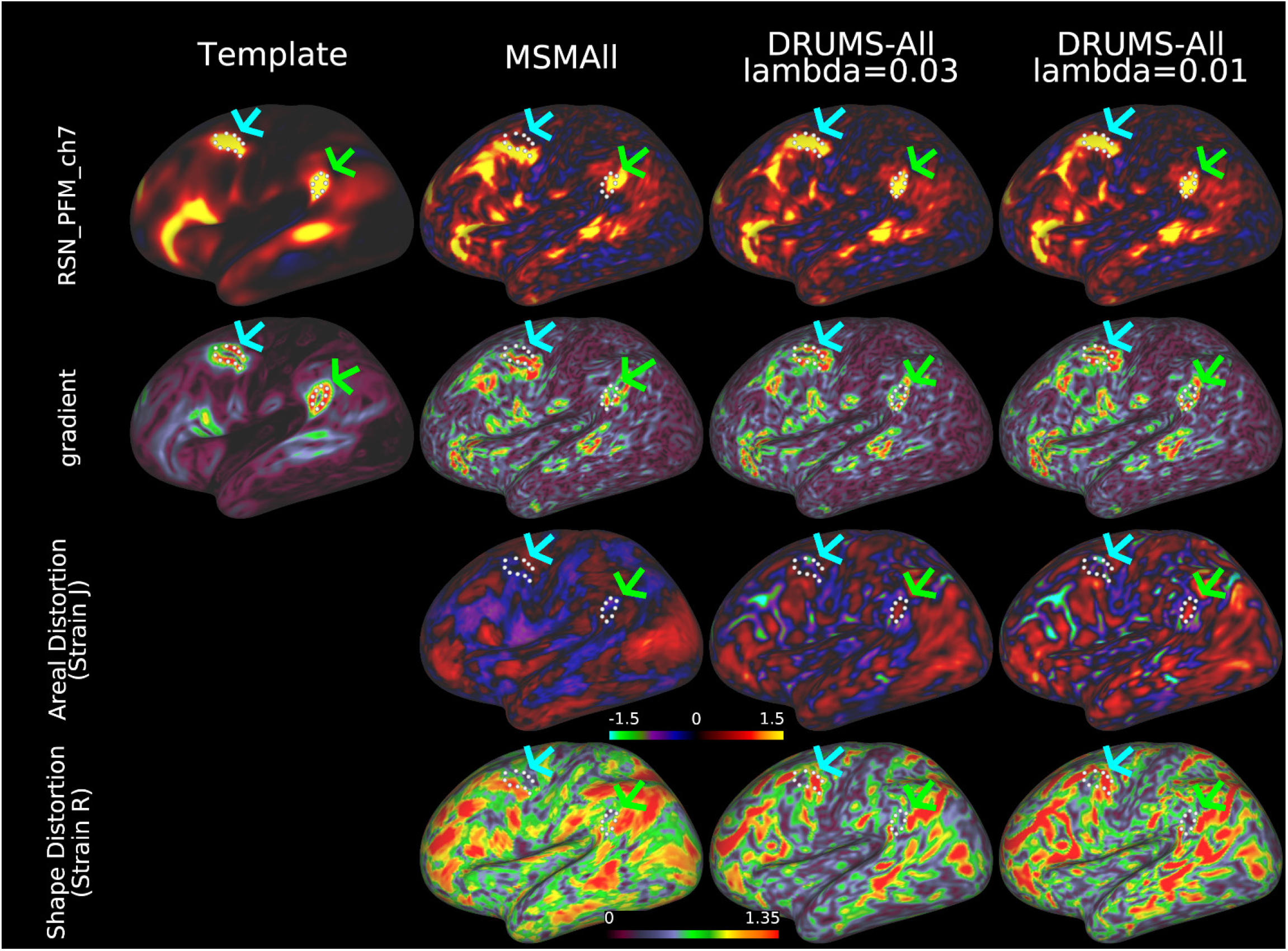
Comparison between MSMAll and DRUMS-All on an example individual. The first row shows an example of a PFM component derived from resting state fMRI, the second row shows the gradient of that component. The lower two rows show the distortion of the registration. https://balsa.wustl.edu/nrG46

In Supplementary Figure 3, we present the same PFM components as Supplementary Figure 2, another subject is presented for better illustration. In this example, DRUMS-SupSulc improves the alignment of the RSN components, which is shown by increased overlap between the two marked regions and the template. By incorporating multimodal feature maps, DRUMS-SupSulc enhances alignment quality in some regions.

#### 8.5. Figures for the right hemisphere

**Supplementary Figure 3:**
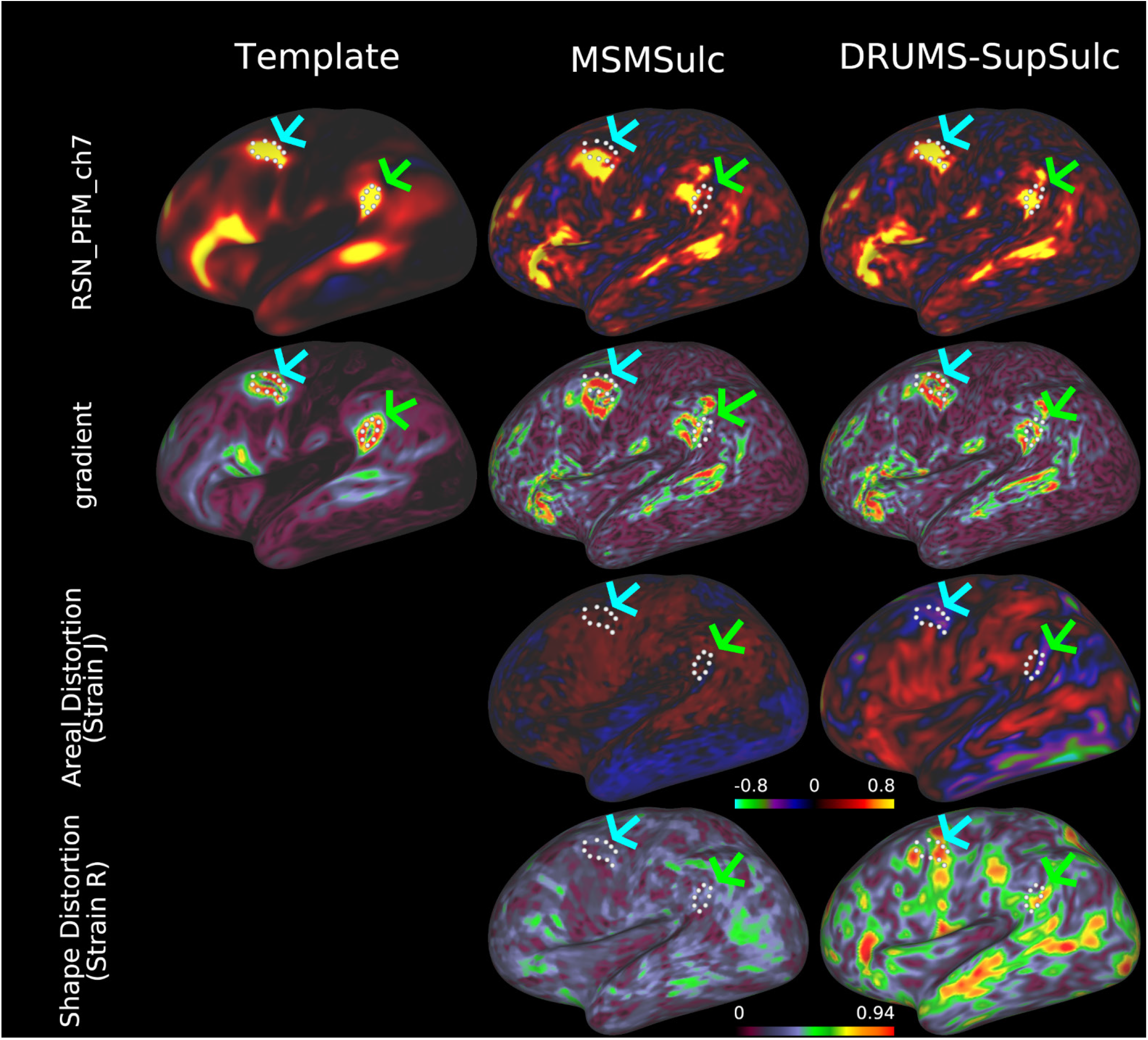
Comparison between MSMSulc and DRUMS-SupSulc on an example individual. The first row shows an example of a PFM component derived from resting state fMRI, the second row shows the gradient of that component. The lower two rows show the distortion of the registration. https://balsa.wustl.edu/gGgZ1

**Supplementary Figure 4:**
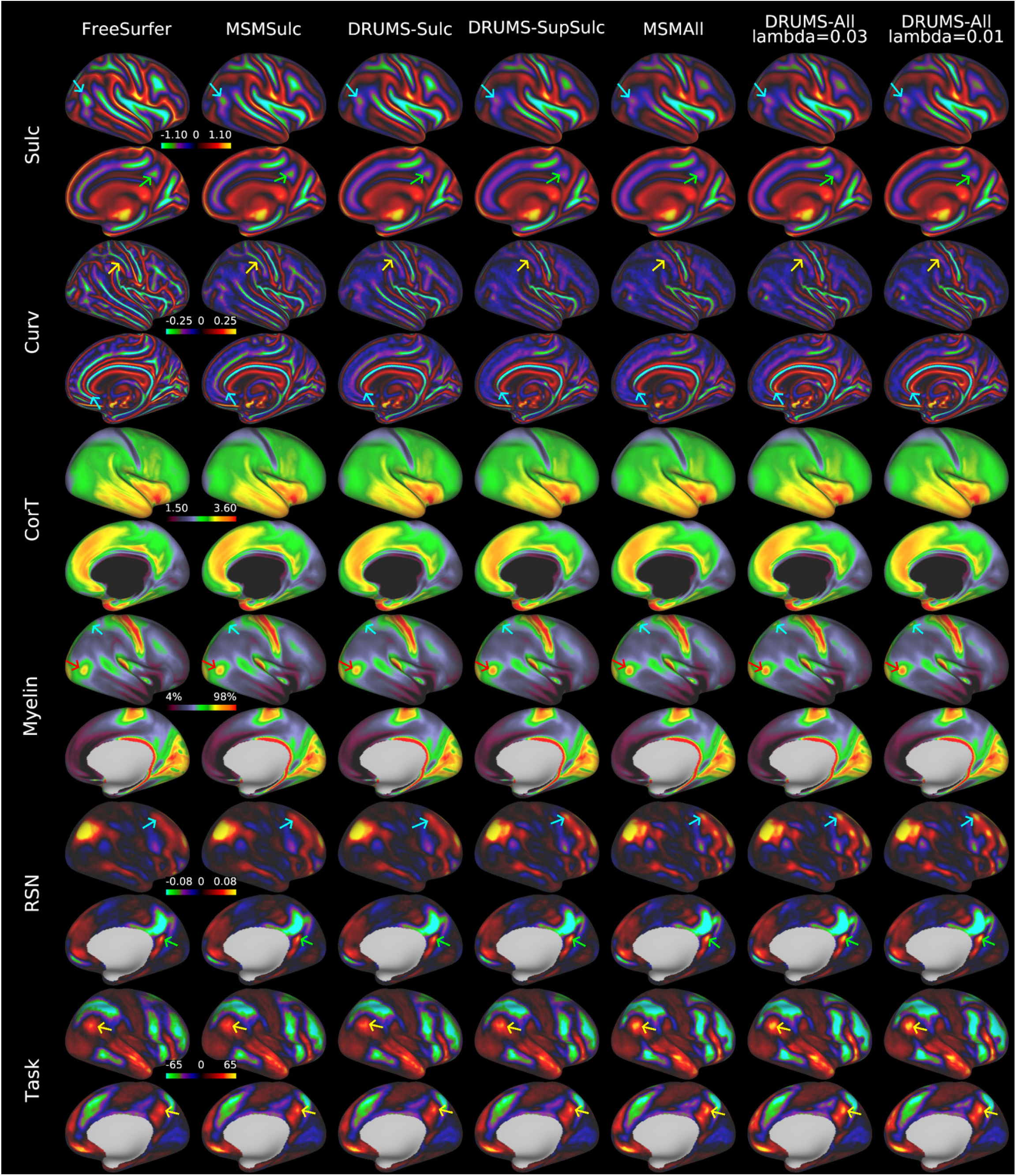
Group average across 475 subjects in the testing set. Each two rows show lateral and medial view of the right hemisphere. Each column shows a registration. https://balsa.wustl.edu/3GqD4

**Supplementary Figure 5:**
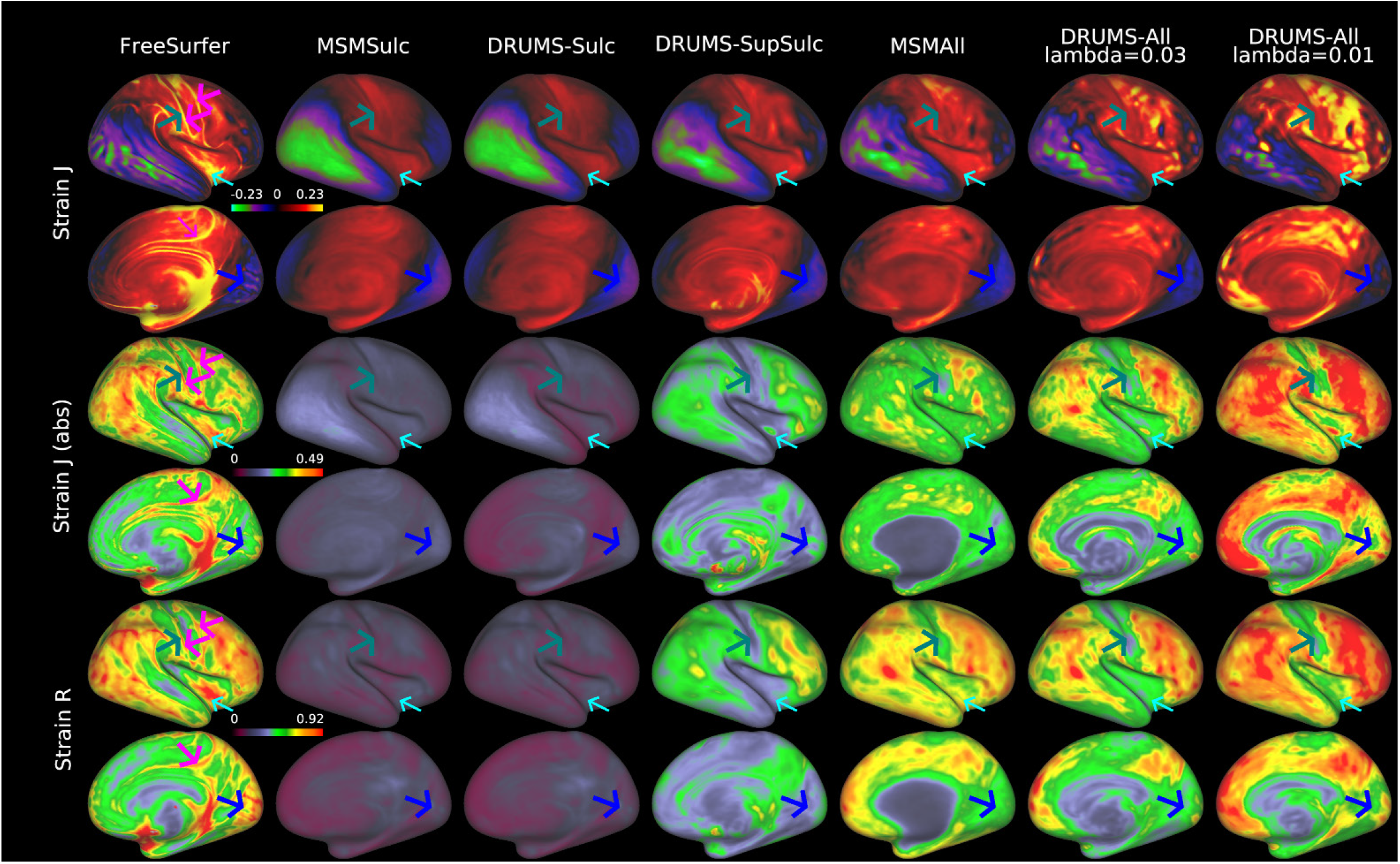
Group average distortion for the right hemisphere across 475 subjects in the testing set. Strain J and R values are log_2_ based. https://balsa.wustl.edu/K82rk

**Supplementary Figure 6:**
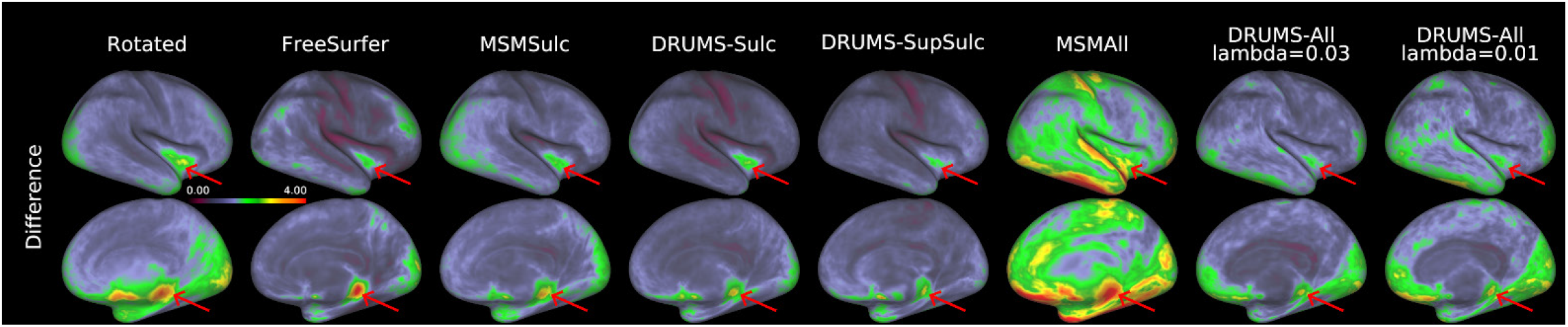
Differences in vertex coordinates between test–retest subject pairs for each registration method for the right hemisphere. Higher values indicate larger differences in vertex coordinates. https://balsa.wustl.edu/x4DMK

**Supplementary Figure 7:**
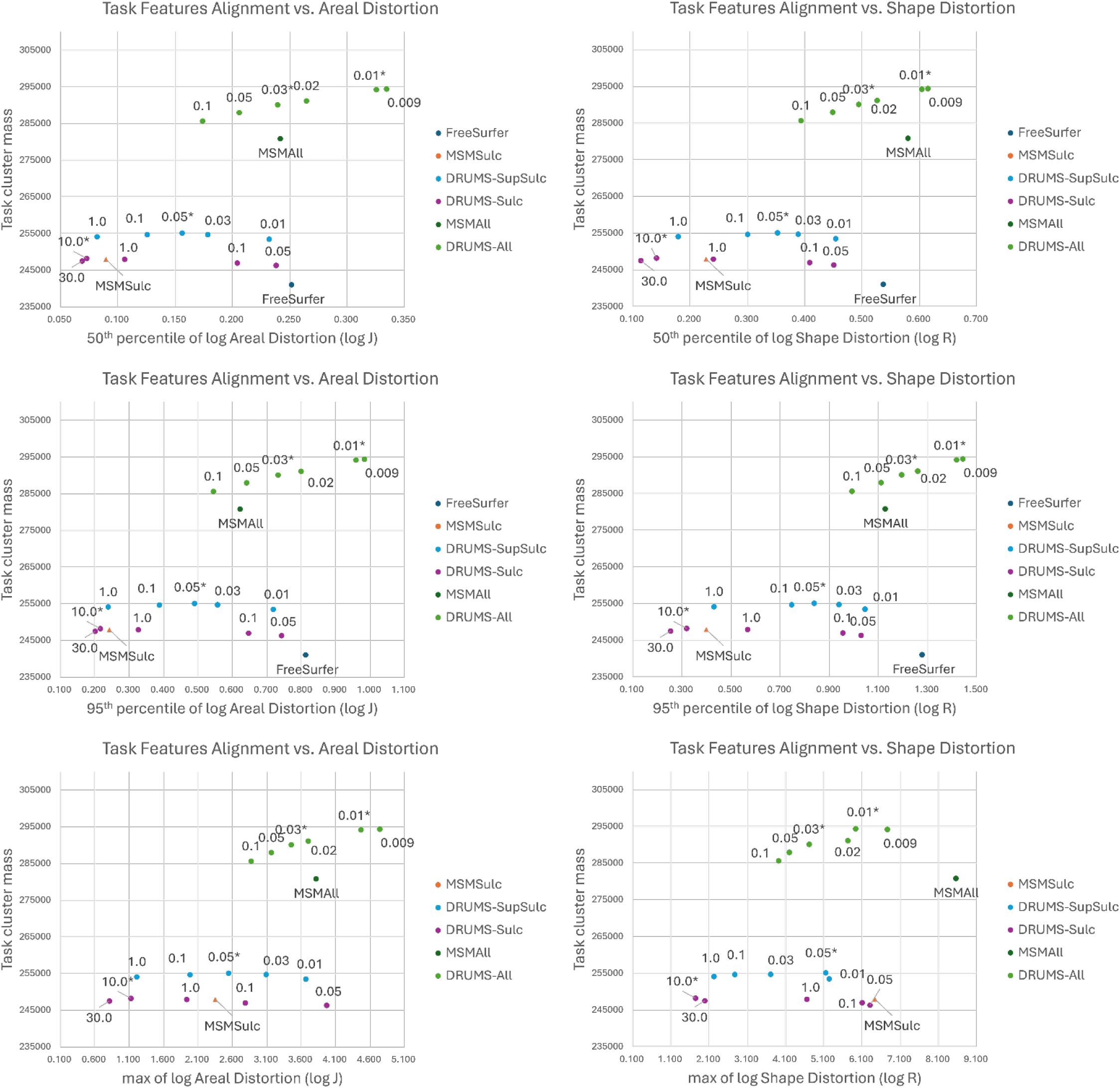
Relationship between task cluster mass and areal distortion. The x-axes show the amount of areal distortion (left three) and shape distortion (right three) with log_2_ transformation, and the y-axes show the task cluster mass. The triangles represent folding based registration; the points represent registration based on functional features. The lambda value used to regularize each model is labeled next to the data point (lower lambda allows more distortion). Notice that the lambda values are not comparable across different models due to their difference in loss functions. The results of models with lambda marked with * are reported in other tables and figures. Datapoints for Freesurfer are removed from both figures for maximum distortions.

